# Spherical deconvolution with tissue-specific response functions and multi-shell diffusion MRI to estimate multiple fiber orientation distributions (mFODs)

**DOI:** 10.1101/739136

**Authors:** Alberto De Luca, Fenghua Guo, Martijn Froeling, Alexander Leemans

## Abstract

In diffusion MRI, spherical deconvolution approaches can estimate local white matter (WM) fiber orientation distributions (FOD) which can be used to produce fiber tractography reconstructions. The applicability of spherical deconvolution to grey matter (GM), however, is still limited, despite its critical role as start/endpoint of WM fiber pathways. The advent of multi-shell diffusion MRI data offers additional contrast to model the GM signal but, to date, only isotropic models have been applied to GM. Evidence from both histology and high-resolution diffusion MRI studies suggests a marked anisotropic character of the diffusion process in GM, which could be exploited to improve the description of the cortical organization. In this study, we investigated whether performing spherical deconvolution with tissue specific models of both WM and GM can improve the characterization of the latter while retaining state-of-the-art performances in WM. To this end, we developed a framework able to simultaneously accommodate multiple anisotropic response functions to estimate multiple, tissue-specific, fiber orientation distributions (mFODs). As proof of principle, we used the diffusion kurtosis imaging model to represent the WM signal, and the neurite orientation dispersion and density imaging (NODDI) model to represent the GM signal. The feasibility of the proposed approach is shown with numerical simulations and with data from the Human Connectome Project (HCP). The performance of our method is compared to the current state of the art, multi-shell constrained spherical deconvolution (MSCSD). The simulations show that with our new method we can accurately estimate a mixture of two FODs at SNR≥50. With HCP data, the proposed method was able to reconstruct both tangentially and radially oriented FODs in GM, and performed comparably well to MSCSD in computing FODs in WM. When performing fiber tractography, the trajectories reconstructed with mFODs reached the cortex with more spatial continuity and for a longer distance as compared to MSCSD and allowed to reconstruct short trajectories tangential to the cortical folding. In conclusion, we demonstrated that our proposed method allows to perform spherical deconvolution of multiple anisotropic response functions, specifically improving the performances of spherical deconvolution in GM tissue.

**Highlights:**

- We introduce a novel framework to perform spherical deconvolution with multiple anisotropic response functions (mFOD)
- We show that the proposed framework can be used to improve the FOD estimation in the cortical grey matter
- Fiber tractography performed with mFOD reaches the cortical GM with more coverage and contiguity than with previous methods
- The proposed framework is a first step towards GM to GM fiber tractography

## Introduction

Diffusion Magnetic Resonance Imaging (dMRI) is a non-invasive technique sensitive to the microscopic motion (diffusion process) of water molecules^1^. In biologic tissues, the diffusion process is influenced by the presence of biologic membranes and macromolecules^2^, which can hinder and/or restrict the molecular random walk in both isotropic and anisotropic fashions. In 1994, Basser et al.^3^ proposed the diffusion tensor imaging (DTI) framework to characterize the anisotropy of hindered diffusion processes. Later, it was shown that when the diffusion process exhibits a sufficient degree of anisotropy, its principal direction can be computed^3^ and eventually tracked^4^, ultimately leading to what is nowadays known as fiber tractography^5,6^ of the brain white matter (WM).

DTI-based fiber tractography is efficient in terms of acquisition and computation but suffers from a number of limitations. Firstly, the technique fails when multiple brain pathways coexist in the same spatial location^7^, which has been estimated to be the case in over 90% of the human WM^8^ and which is not likely to be solved by technological improvements as higher imaging resolutions^9^. Secondly, DTI-based fiber tractography only applies to data acquired within the Gaussian diffusion regime, and thus cannot take advantage of the strong diffusion weightings which are typical for high angular resolution diffusion imaging methods^10^. Due to these considerations, DTI is not likely the method of choice to study a heterogeneous structure as grey matter (GM), which is characterized by low fractional anisotropy (FA) and, hence, high uncertainty of the principal diffusion direction.

Spherical deconvolution^11,12^ is one of the most popular methods to reconstruct fiber orientation distributions (FOD) and to perform fiber tractography in WM. Conversely, fiber tractography based on spherical deconvolution methods has found very limited application to investigate the organization of the cortical GM, despite its pivotal role as endpoint of most axonal bundles. This is most likely due to three unsolved challenges which limit the performance of spherical deconvolution in GM. Firstly, the presence of superficial WM next to the inner cortical surface limits the transition of the streamlines from the deep WM to the cortical GM, and vice versa^13^. Secondly, the fiber orientations of the superficial WM are typically tangent to the cortical surface, causing the premature termination of streamlines to and from the cortical banks and sulci, which results in the well-known gyral bias towards the cortical crowns^14^. Thirdly, the FODs derived in the cortex are noisy and exhibit spurious peaks, preventing reliable propagation of the streamlines in the cortical ribbon. Importantly, these limitations in fiber tractography propagate to other analysis techniques making use of fiber reconstructions as, among others, structural connectivity approaches^15^, cytorachitectonical analyses^16^, and pre-surgical planning^17^. In this work, we focus on the third challenge, and propose a novel framework to improve the reliability of the fiber orientations determined with spherical deconvolution in the cortical grey matter. Of note, previous works have proposed geometric heuristics to determine the fiber orientation in the cortex based on the cortex normal^18^ or on the vector field from the inner to the outer cortex^19^. Although these methods have the merit to work with commonly available datasets, they oversimplify the way axons traverse the cortex^20^, and do not allow to take into account the simultaneous presence of tangential and perpendicular fiber orientations in the cortex^20,21^, which is region- and layer-specific.

The advent of specialized techniques for diffusion imaging, such as ultra-strong gradients^22^ and simultaneous multi-slice^23^, are rapidly increasing the imaging quality and resolution achievable in dMRI. This allows to appreciate new details of the human cortex, as recently reviewed by Assaf^24^. In ex-vivo studies^20,21,25^, it has been consistently shown that the diffusion process observed in the cortex has a large anisotropic component perpendicular to the cortical folding, but also that tangential directions can be observed, for instance, in the central sulcus^26^. In particular, the anisotropy of the cortex has been observed with ex-vivo MRI in animals, as well as in the human brain, both ex-vivo^21,27,28^ and in-vivo^26,29^. Such high-quality data acquisitions may soon be routinely available for in-vivo applications, offering unprecedented research possibilities^30^ in fiber tractography of GM, but a dedicated framework has not yet been investigated.

Multi-tissue spherical deconvolution approaches^31^ have been suggested to further increase the accuracy of FOD reconstructions in WM by accounting for partial volume effects, e.g. in spatial locations where WM overlaps with GM or cerebrospinal fluid (CSF). Although GM and CSF are merely considered nuisance factors^32^, such framework highlights the feasibility of explicitly accounting for multiple tissues in spherical deconvolution approaches. By separating multiple partial volume effects, multi-tissue approaches have shown improved FOD reconstructions in adults^31^, neonates^33^ and in patients with Parkinson’s disease^34^. In current state-of-the-art multi-shell spherical deconvolution, only WM is assumed to be anisotropic, while GM is regarded as an isotropic tissue. To estimate the WM FOD, a WM fiber model – or response function – is estimated from voxels with a single fiber population, which are typically located in large bundles^35^ such as the corpus callosum or the corticospinal tract. The response function is then assumed to be valid throughout the brain. While the validity of this representation has been questioned even in WM itself^36^, it is highly unlikely that it can adequately represent the anisotropic signature of the cortex, which in addition to axons also contains dendrites and cell-bodies^37–40^.

In this study, we investigate whether the spherical deconvolution approach can be revisited to consider the different signal characteristics of WM and GM by explicitly modelling both signal mixtures simultaneously. This novel framework allows to reconstruct multiple FODs (mFOD) and is ultimately aimed at improving the FOD estimation and fiber tractography of GM while retaining state-of-the-art performances in WM. In our proposed method, we disregard the existence of a single response function for the brain, and investigate whether it is feasible to simultaneously estimate multiple anisotropic FODs corresponding to tissue-specific response functions. We also evaluate whether this improves the fiber orientation characterization in GM, and fiber tractography in GM as compared to existing spherical deconvolution methods. The mFOD method is in principle also applicable to deep GM structures such as the thalamus, which is often of interest in fiber tractography applications due to its hub role in several fiber bundles^41^. In this work, however, we focus on the study on the cortical GM and only show proof-of-concept of the applicability of mFOD to deep GM, which should be investigated in more detail in future work. Preliminary results of this work have been presented at the 2019 ISMRM meeting in Montreal, Canada^42^.

## Theory and methods

The following paragraphs present the mFOD framework and explain the qualitative and quantitative analyses that we performed using both simulations and in-vivo data.

### Generalized Richardson Lucy (GRL)

In previous work^43,44^, we introduced a multi-shell spherical deconvolution framework named Generalized Richardson Lucy (GRL) to account for multiple tissue types, such as WM, GM and cerebrospinal-fluid (CSF), as shown in Eq. 1.

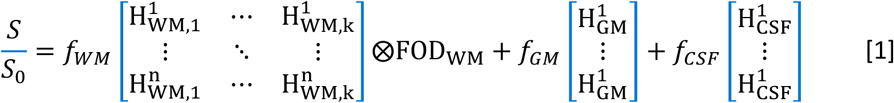

In Eq. [1], *S* is the signal collected at *n* diffusion weightings, *S*_*0*_ is the non-diffusion-weighted signal, the H matrices represent the signal model associated to WM, GM and CSF, *f* is the associated signal fraction, and FOD_WM_ is the fiber orientation distribution associated to WM. The columns of the deconvolution matrix shown in Eq. [1] represent the possible solution of the spherical deconvolution on the unit sphere, resulting in a total number of *k*+2 columns, as GM and CSF are both considered isotropic.

### Multiple fiber orientation distributions (mFOD)

In the mFOD framework, we redesigned the deconvolution matrix to allow for an arbitrary number of independent anisotropic deconvolution models. Considering solving the spherical deconvolution problem once more for WM, GM and CSF, but now accounting for anisotropic diffusion in both WM and GM, we can write:

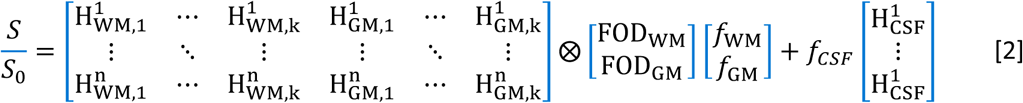

Eq. [2] can be solved using the iterative least-squares minimizer previously introduced for GRL but replacing the Richardson-Lucy deconvolution scheme with a regularized non-negative least-squares solver^45^, producing multiple independent FODs.

### Signal models (response function)

To underline the model independence of the mFOD framework, the H-matrices of WM, GM and CSF, shown in Eq. 2, were evaluated with three different models. The columns of the H-matrices are generated with instances of a specific model, e.g. the signals are generated with a predefined set of parameters and then projected along a number of spatial directions defined on the unit sphere. In other works, the H-matrix is referred to as deconvolution matrix, or response function or kernel. H_CSF_ was generated using the ADC model with the typical diffusion coefficient of free water at 37 degrees (3.0 × 10^−3^ mm2/s). The columns of H_WM_ and H_GM_ were generated using the DKI model^46^ with isotropic kurtosis^47^, and the NODDI model^38^, respectively, both in simulations and with in-vivo data. For both models, the deconvolution matrix included 300 possible solutions equally distributed on the unit-sphere, which results in an angular resolution of about 4°. The DKI model was chosen as it can represent the anisotropy of the WM signal while also considering non-Gaussian diffusion effects. The NODDI model has been shown to characterize the GM microstructure better than tensor models^48^, and allows to take into account the larger presence of cell bodies in GM as compared to WM. It should be noted that mFOD is a generic framework that could be used in combination with models other than DKI and NODDI, such as geometric models^49^ or data driven representations^50^. The combination of the simplified DKI and NODDI models should thus not be regarded necessarily as the optimal choice, but rather as a proof of concept. The parameters of the models were determined as explained in the following paragraphs.

### Simulations

The feasibility of simultaneously computing multiple anisotropic FODs corresponding to different models was evaluated numerically. H_WM_ was generated with the DKI model by setting the eigenvalues of the diffusion tensor as (1.7, 0.3, 0.3) x 10^−3^ mm^2^/s, and isotropic kurtosis equal to 0.6. H_GM_ was generated with the NODDI model by simulating values of intra to extra-cellular ratio 0.4, parallel diffusivity 1.7×10^−3^ mm^2^/s, concentration parameter of the Watson distribution equal to 1.0, and no partial volume of free-water was considered as the mFOD approach explicitly accounts for it. The parameters used for generating H_WM_ and H_GM_ matched the average values observed with in-vivo data.

Two sets of simulations were generated using the 3-shells HCP acquisition protocol. Simulation I aimed at evaluating the robustness to noise of the mFOD approach, whereas simulation II characterized its ability to separate partial volume effects for a given SNR level.

### Simulation I

To investigate the feasibility of mFOD, we generated three sets of partial volume effects of WM and GM, to simulate the transition from deep WM to GM. The first configuration simulates the partial volume of 1 WM-like fiber and CSF, which in-vivo can be observed in the deep WM, for instance in proximity of the ventricles. In this simulation, f_WM_ was set equal to 0.8, and f_GM_ equal to 0.2. The second configuration simulates the partial volume of 1 GM-like fiber and CSF with signal fractions f_GM_=0.8, f_CSF_=0.2, and is meant to reproduce the setting observed in-vivo in the outer cortex. The third configuration simulates the partial volume between 2 WM-like fibers with crossing angle 45°, and 1 GM-like fiber crossing the first WM-like fiber with angles 30°-60°-90°, respectively, to mimic the transition of a WM fiber to GM through the superficial WM. For this configuration, the signal fractions f_WM_ and f_GM_ were both set equal to 0.5. The fourth configuration simulates the crossing of a WM-like and a GM-like fiber, which simplifies what may be observed in-vivo after the superficial WM is crossed. In this case, the simulation was repeated for crossing angles in the range 10° - 90° with signal fractions f_WM_ and f_GM_ both equal to 0.5. The fifth configuration is representative of a voxel containing mostly GM. For this configuration, we used the same settings of the previous simulation but adjusted the values of the signal fractions to f_WM_ = 0.2 and f_GM_ = 0.8. The effect of noise was simulated by adding 1000 Rician realizations in correspondence of SNR levels 10, 20, 30, 40, 50, 60, 70, and 150. For each SNR level, we evaluated the error between the estimated and the simulated signal fractions, as well as the angular error between the simulated and the estimated FOD directions.

Additional simulations were performed to understand the performance of mFOD as function of the employed signal model and of the acquisition protocol and can be found in the Supplementary Material.

### Simulation II

Three partial volume configurations were generated in analogy with simulation I – configuration V with a crossing angle equal to 75°, but increasing the ratio between the two simulated tissues from 0 to 1 with step 0.1. Rician noise was added by considering 1000 realizations at SNR 50. The error between the simulated and the estimated signal fractions was determined.

### In-vivo data processing

Two subjects from the pre-processed dataset of the Human Connectome Project (HCP)^22,51^ were randomly chosen and included in this study. The datasets included a T1-weighted image acquired at a resolution of 0.7 mm isotropic, and a dMRI dataset acquired at 1.25 mm isotropic. The diffusion datasets consisted of 18 b = 0 s/mm^2^ volumes in addition to 270 volumes acquired by sampling diffusion weightings b = 1000, 2000, and 3000 s/mm^2^ along 90 directions each. One of the two subjects was chosen to showcase the main results of the mFOD approach, using the native dMRI space (1.25 mm) as analysis space.

Generalizability of the methods was then shown on the second subject, using the T1 resolution (0.7 mm) to highlight some features of the mFOD method. The T1-weighted data of the first subject was rigidly registered to the pre-processed dMRI data. Then, the tissue type segmentations of both datasets in WM, GM and CSF were derived with FSL FAST^52^. The SNR of the two datasets was determined by computing the average and the standard deviation of the signals at b = 0 s/mm^2^ in two regions of interest manually placed in deep WM and in the parietal cortex, respectively. The average SNR at b = 0 s/mm^2^ of the first dataset was 20 ± 4 in WM and 32 ± 9 in GM. The average SNR at b = 0 s/mm^2^ of the second dataset was 20 ± 5 in WM and 27 ± 8 in GM.

The dMRI data was fit with both the “Multi-Shell Constrained Spherical Deconvolution” (MSCSD)^31^ (only data at 1.25 mm) and the mFOD approaches using custom implementations in MATLAB R2018b (The Mathworks Inc.) and functions from the ExploreDTI toolbox^53^. The code of mFOD is freely available online as part of the “MRIToolkit” toolbox for MATLAB (https://github.com/delucaal/MRIToolkit). MSCSD was initialized as previously suggested independently on each dataset with spherical harmonics of order 8 by using the tissue maps derived from the T1-weighted image. For the generation of the H-matrices used for the mFOD approach, a whole brain DKI and NODDI fit were performed with the dMRI data of the two HCP datasets at 1.25 mm resolution. For the WM H-matrix, the eigenvalues of the diffusion tensor and the isotropic kurtosis were averaged within a white matter mask defined by fractional anisotropy values above 0.7. This resulted in eigenvalues [1.3, 0.5, 0.5] x 10^−3^ mm^2^/s and isotropic kurtosis value 0.64. For the H-matrix of GM, the NODDI metrics were averaged within the GM segmentation obtained with the T1-weighted image, which resulted in intra to extra-cellular ratio 0.4, parallel diffusivity 1.7×10^−3^mm^2^/s and Watson’s concentration parameter (*κ*) equal to 1. In this work, identical H-matrices were used for the mFOD analysis of both datasets as proof-of-concept. More generally, however, we recommend optimizing the signal models on a per-subject basis, especially when studying subjects with large lifespan or in patients. A graphical representation of the H-matrices (response functions) used for the analysis of in-vivo data is shown in Figure 1 for the data shell at b = 3000 s/mm^2^. The figure shows that the H-matrix corresponding to mFOD_1_ is pancake-like and appears similar to the response function commonly used in constrained spherical deconvolution. The NODDI-based H-matrix used for mFOD_2_ is also clearly anisotropic and assumes a zeppelin-like shape.

**Figure 1:**
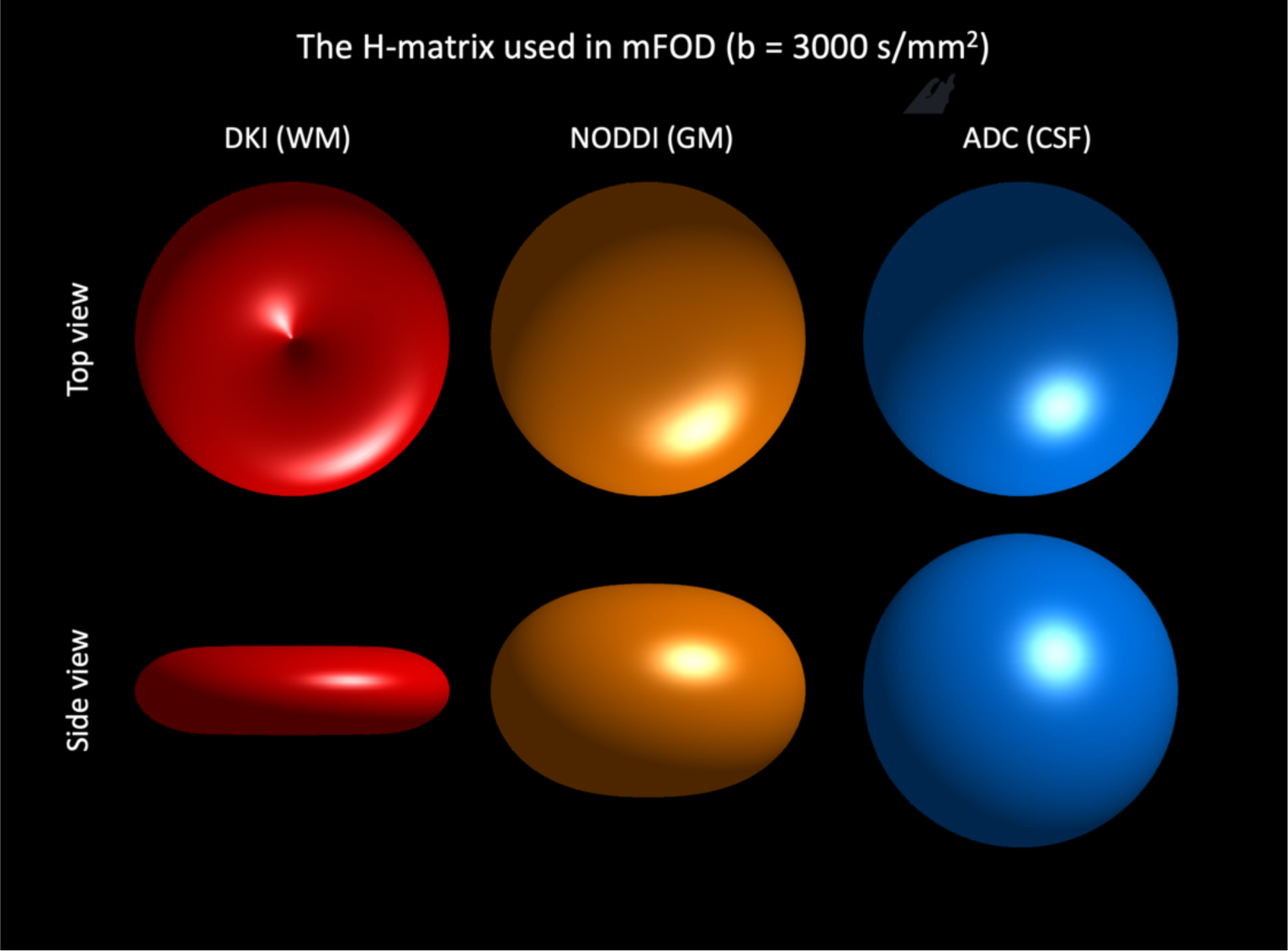
A graphical representation of the H-matrices (response functions) used in mFOD in correspondence of data at b = 3000 s/mm^2^. The white matter (WM) signal is represented with the diffusion kurtosis imaging (DKI) model and shows a highly anisotropic profile. The NODDI model is used to represent the anisotropic diffusion signal in grey matter (GM). The corresponding H-matrix results in an anisotropic – zeppelin-like – 3D profile.

### FODs and signal fractions

The MSCSD fit produced signal fraction maps associated to WM, GM and CSF, and one FOD describing WM orientations. The fit of the mFOD approach resulted in the same number of fractional maps but two FODs, mFOD_1_ describing WM geometry, and mFOD_2_ describing GM geometry, respectively.

The signal fraction maps computed with MSCSD and mFOD were quantitatively compared by evaluating their Pearson correlations. To this end, f_WM_ and f_GM_ were evaluated, respectively, in WM and GM masks derived from the T1-weighted image. Next, the signal fraction maps from both mFOD and MSCSD were visually compared to those obtained from the T1-weighted volume. Further, the FODs computed with MSCSD and the mFOD framework were compared in selected areas of the cortex and of deep GM, to determine whether mFOD_2_ provided additional information in GM as compared to mFOD_1_ and MSCSD.

The FODs computed with the mFOD method can be either analyzed separately, or merged to combine accurate geometrical information in both WM and GM. As a proof of concept, we combined the two FODs by considering their normalized sum weighted by the respective signal fractions (mFOD-WS).

One of the difficulties when validating in-vivo dMRI techniques is the lack of ground truth. MSCSD can be regarded as a gold standard in WM, but debatably not in GM. In a first analysis, we investigated to which extent the principal diffusion directions of the mFOD-WS agree with those of MSCSD in WM, GM and at the WM/GM interface. A voxel was assigned to the latter class when its probability of belonging to both WM and GM was non-zero in the T1-weighted segmentation. Per voxel, up to three peaks were identified for both FODs, then sorted per descending amplitude values. Subsequently, the dyadic angle between each mFOD-WS peak and the closest MSCSD peak was determined.

If discrepancies are observed in GM, it remains impossible to ultimately judge which method is the most accurate, but comparisons with a third method could provide additional interpretations. Anatomical knowledge as well as previous MRI studies^30^ suggest that the dominant diffusion directions in GM should be perpendicular to the cortical folding, with exceptions such as the primary visual cortex^20^. We therefore derived the normal vectors to the cortical folding based on the structural tensor^54^. In short, a 3D volume was initialized with 1-values in correspondence of the GM/WM and GM/CSF interfaces, as derived from the T1-weighted segmentation. The linear interpolation field between the two surfaces was derived, and the corresponding vectors computed by computing the X-Y-Z first order derivatives of the volume. Subsequently, we determined the minimum dyadic angle between the normal vectors and any peak orientation determined with the MSCSD and mFOD-WS results, respectively.

### Fiber tractography

#### Tracking parameters

We investigated fiber tractography results obtained with MSCSD, mFOD_1_, mFOD_2_ and mFOD-WS, with a particular focus on GM areas. Deterministic fiber tractography was performed in *ExploreDTI* by using identical parameters for all methods: FOD threshold 0.1 (default), minimum/maximum fiber length equal to 20/500mm, angle threshold 45 degrees, and step-size 0.6 mm. Fiber tractography was also performed in an axial region of interest overlapping the central sulcus in a single slice, and using FOD thresholds of 0.1 and 0.3, minimum fiber lengths of 10 and 50 mm, and step-size of 0.35 mm.

Firstly, FODs computed with the MSCSD and mFOD-WS approaches were computed in a frontal region and color encoded with a custom colormap assigning red, yellow and blue to pathways traversing WM, GM and CSF, respectively. Subsequently, whole-brain tractography of all FODs was performed by seeding all voxels with an FOD value above the default threshold. The fiber bundles were color encoded both with the conventional RGB color scheme^55^ and the above-mentioned tissue-type color scheme, to convey both directional information and specificity of the FODs to the different tissue types.

Fiber pathways obtained with MSCSD and mFOD in a 3.75 mm wide sagittal slab were color encoded by normalized distance between the GM/WM interface and the WM/CSF interface – as obtained from the T1-weighted segmentation – and also directly overlaid, to appreciate differences in GM tracking distance and density between the compared methods.

Subsequently, we evaluated the number of tract pathways generated with MSCSD and mFOD-WS, which is informative given that the seed-points were kept constant and that a deterministic tracking algorithm was used. Additionally, we counted the fraction of plausible tract pathways, i.e., trajectories having at least one endpoint in grey matter, and the spatial coverage of tractography within GM when considering only start/endpoints of tracks. To qualitatively evaluate whether mFOD-WS could benefit fiber tractography of deep grey matter regions, we performed deterministic tractography of both mFOD-WS and MSCSD in an axial sub-region of the thalamus.

To qualitatively assess whether the mFOD framework can reconstruct previously reported fibers tangential to the cortex^26^, we investigated the fiber tractography results with the mFOD-WS approach.

Fiber tractography was seeded in a single axial slice, then compared to the cortical surface reconstructed with the CAT12 toolbox^56^ for MATLAB.

## Results

### Simulations

Fig. 2 and Fig. 3 show the signal fractions estimated with the mFOD approach in simulation I, where different mixtures of WM, GM and CSF simulate 5 configurations at multiple SNR Levels. Fig. 2 highlights the results of configurations I, II, and V with crossing-angle 90°, which are the main focus of this work. The figure shows that at SNR level equal or greater than 20, the mFOD method disentangles mixtures of WM and CSF (configuration I) with signal fraction errors below 2.2% and angular errors below 3 degrees. The separation of GM from both CSF (configuration II) and WM (configuration V) is more challenging and requires higher SNR to achieve an error below 10%. In configuration II, the signal fraction errors at SNR 50 are about 9%, 11% and 2%, for WM, GM and CSF, respectively, and the angular error on the GM FOD is 8 degrees. In configuration V, when WM and GM-like fibers cross with angle equal to 90°, the corresponding signal fraction errors at SNR 50 are 7%, 10% and 2%, and the angular errors on the WM and GM FODs are 4 and 11 degrees, respectively.

**Figure 2:**
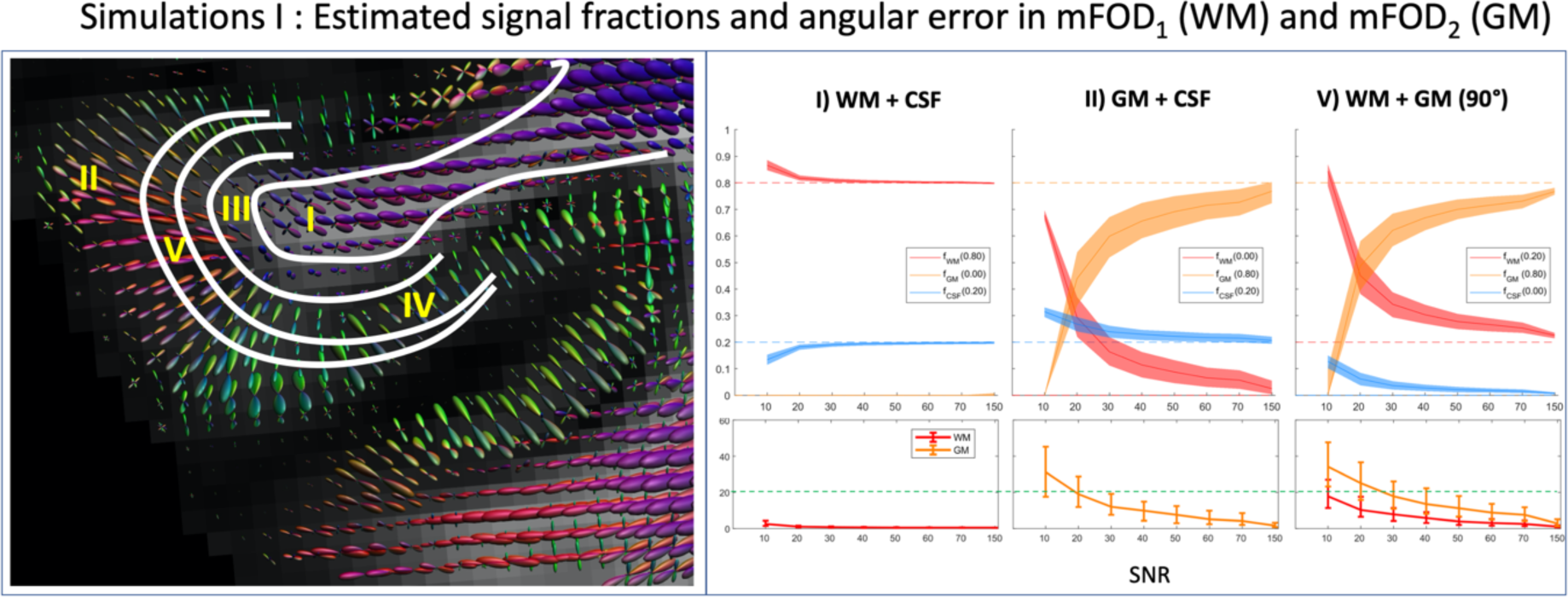
Results of simulation I. Five configurations of WM and GM mixtures simulating I) a WM-like fiber with CSF partial volume (20%), II) a GM-like fiber with CSF partial volume (20%), III) two crossing WM-like fibers and a GM-like fiber, IV) a WM-like and a GM-like crossing fibers with equal volume fraction, and V) a WM-like and a GM-like crossing fibers with volume ratios 0.2/0.8, respectively. The simulations are designed to reproduce some physiologically plausible configurations, as schematically shown in the left sketch of a cortical gyrus. For each simulation, the top plot shows the 25^th^ – 75^th^ percentile (shaded area) and the median (solid line) of the signal fractions estimated with the mFOD approach for different SNR levels The bottom plot shows the angular error of the main direction of the WM and GM FODs as compared to the ground truth value. The solid line represents the median value, whereas the error bars the 25^th^ and 75^th^ percentile. The simulations were performed using anisotropic response functions for WM and GM, and an isotropic response function for CSF. The green dotted line corresponds to an angular error equal to 20°. To properly detect the main peak orientation associated to GM, a high SNR (≥50) is required.

**Figure 3:**
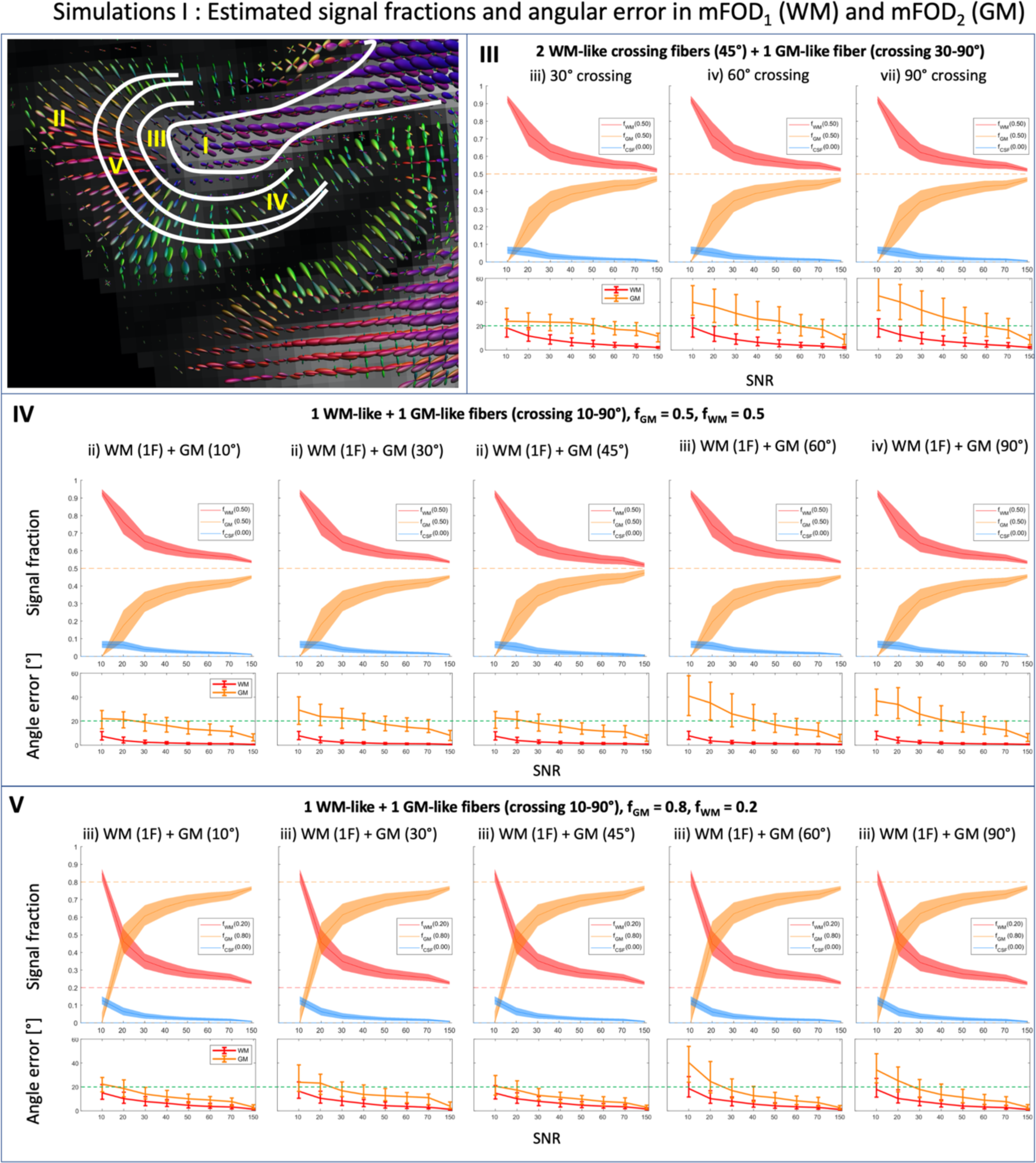
Results of simulation I. Five configurations of WM and GM partial volume simulating I) a WM-like fiber with CSF partial volume (20%), II) a GM-like fiber with CSF partial volume (20%), III) two crossing WM-like fibers and a GM-like fiber, IV) a WM-like and a GM-like crossing fibers with equal volume fraction, and V) a WM-like and a GM-like crossing fibers with volume ratios 0.2/0.8, respectively. For each simulation, the top plot shows the 25^th^ – 75^th^ percentile (shaded area) and the median (solid line) of the signal fractions estimated with the mFOD approach for different SNR levels The bottom plot shows the angular error of the main direction of the WM and GM FODs as compared to the ground truth value. The solid line represents the median value, whereas the error bars the 25^th^ and 75^th^ percentile. The simulations were performed using anisotropic response functions for WM and GM, and an isotropic response function for CSF. The green dotted line corresponds to an angular error equal to 20°. To properly detect the main peak orientation associated to GM, a high SNR (≥50) is required.

Fig. 3 shows the results of simulation I for configurations III-IV-V as function of the crossing angle between WM and GM for SNR levels. In general, the crossing angle between the WM-like and the GM-like fibers seems to marginally affect the signal fraction estimation. At SNR 50, the min-max signal fraction errors of WM, GM and CSF in configuration III are 7% (60°) – 7.1% (30°), 8.8% (45°) – 9.2% (30°) and 1.9% – 1.9%, respectively. The min-max signal fraction errors at SNR 50 for configuration IV are 8.2% (60°) – 8.9 % (10°) for f_WM_, 10.5% (90°) – 11.2% (10°) for f_GM_, 2.2% (90°) – 2.4% (10°) for f_CSF_. For configuration V, the minimum signal fractions error was observed with a 90° crossing configuration, 7.8% for f_WM_, 10.0% for f_GM_ and 2.1% for f_CSF_, whereas the worst error was observed in correspondence of the 10° crossing, 8.4% for f_WM_, 10.6% for f_CSF_ and 2.2% for f_CSF_.

When the SNR level is above 30, the angular error of the WM FOD is below 10° in configuration III and IV independently from the crossing angle, and in configuration V for crossing angles of 45° or above. Achieving a good estimate of the GM FOD at the GM/WM interface is challenging. In configuration III and IV, the angular error of the GM FOD is greater than 10° at SNR < 70, independently from the crossing angle with WM. More reliable estimates of the GM FOD can be achieved in configuration V: at SNR 50, an angular error of about 9.6° is observed for crossing angles of 45° or greater. In Supplementary Figure S1, the simulation of configuration IV with crossing angle 45° was repeated with different values of the NODDI parameter *κ*. The figure shows that at SNR 50 it is possible to estimate the GM FOD with angular errors equal to 8° and 5.6° in correspondence of *κ* = 2 and *κ* = 3, respectively, which are plausible settings in-vivo at the WM/GM interface. The simulation also shows that the angular resolution of the GM FOD improves with larger values of *κ* even if a value *κ* = 1 is used to generate the mFOD signals.

The effect of the acquisition protocol on the performance of mFOD is investigated in Supplementary Figure S2 and S3. Halving the number of gradient directions as compared to the HCP protocol worsens the effective SNR of the data, resulting in larger spread of the signal fraction estimates. The angular error of the WM FOD is minimally affected by the halving of the gradients, whereas slightly worse performances can be observed for the GM FOD. In Supplementary Figure S3, we investigated whether the distribution of the gradients on the outer shell and its diffusion weighting affects the performance of mFOD. Among the tested scenarios, only increasing the maximum diffusion weighting to b = 4000 s/mm^2^ appreciably affects the performance of mFOD, suggesting a better estimation of the GM FOD (−21% angular error at SNR 50) and reduced uncertainty in the signal fractions estimates for larger maximum diffusion weightings.

Fig. 4 shows the signal fractions estimated in simulation II when linearly varying the simulated WM, GM and CSF fractions at SNR 50. For all the three cases, the estimated average signal fractions show excellent agreement with the simulated values. The maximum errors observed for WM, GM and CSF are 13%, 20% and 4%, respectively. The Pearson correlation coefficient between the simulated and estimated signal fractions have values between 0.99 and 1.

**Figure 4:**
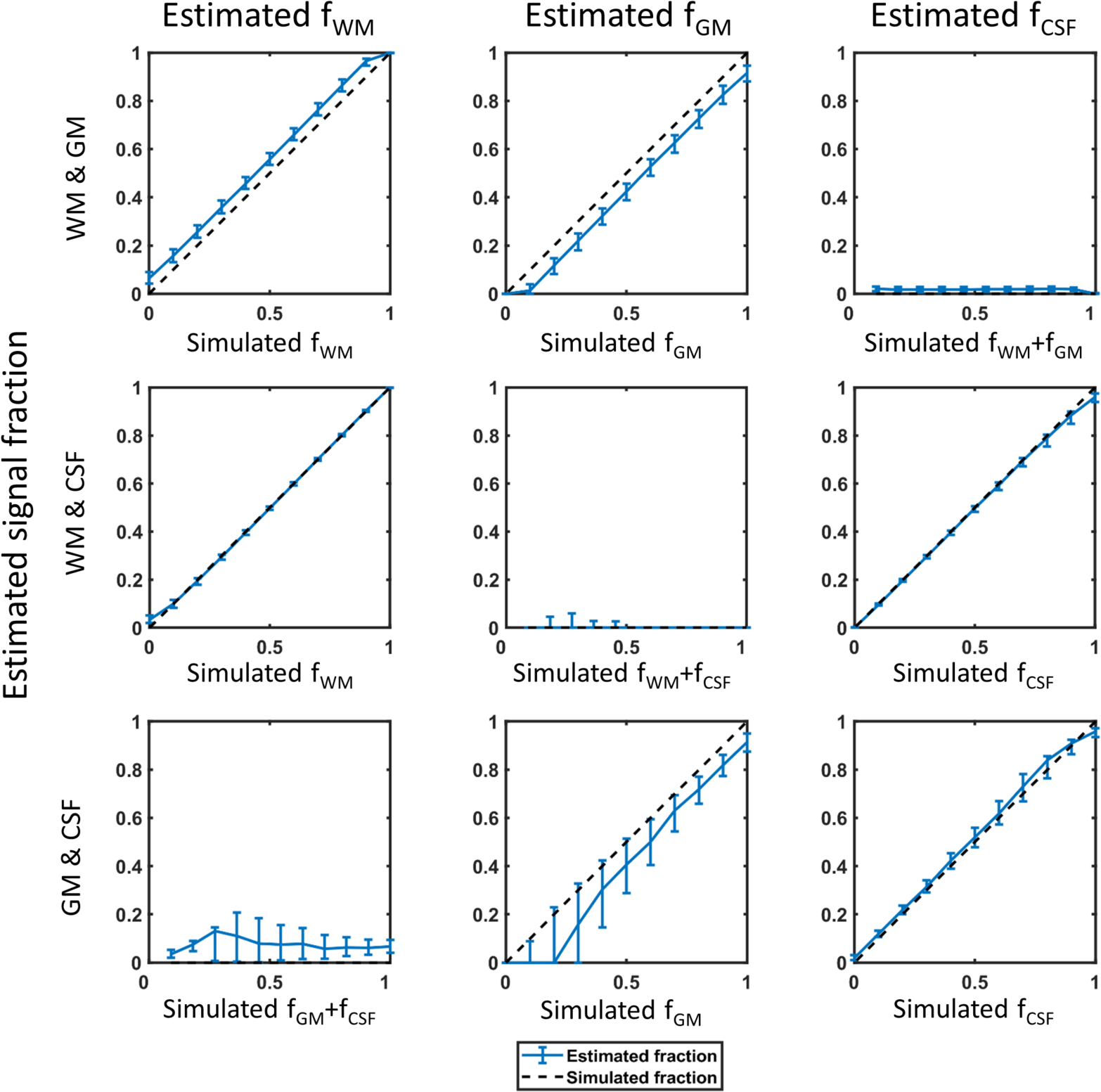
The median value (solid line) and the 25^th^ – 75^th^ percentile (error bars) of the signal fractions estimated with the mFOD approach for the simulated anisotropic WM and GM response functions (columns 1-2) and the isotropic CSF response function (column 3). Different mixtures of the three tissue types were simulated with increasing partial volume and 1000 realizations of Rician noise at SNR 50.

The largest errors were observed when mixing GM and CSF in correspondence of low GM signal fractions.

### HCP data

#### Signal fractions

The WM, GM and CSF signal fractions estimated with the MSCSD and mFOD methods, as well as those derived from the segmentation of the T1-weighted image are shown for an example axial slice in Fig. 5. The maps estimated with the mFOD method exhibit excellent agreement with both MSCSD and the T1-derived segmentation, highlighting the adequacy of the NODDI model and of the chosen parameters to represent the in-vivo signal in GM. The Pearson correlation coefficient between mFOD and MSCSD were 0.96 and 0.95 for f_WM_ and f_GM_, respectively. Notice that the WM fractional maps estimated with both the MSCSD and mFOD methods have non-zero values in some GM areas, in contrast to the T1-derived segmentation.

**Figure 5:**
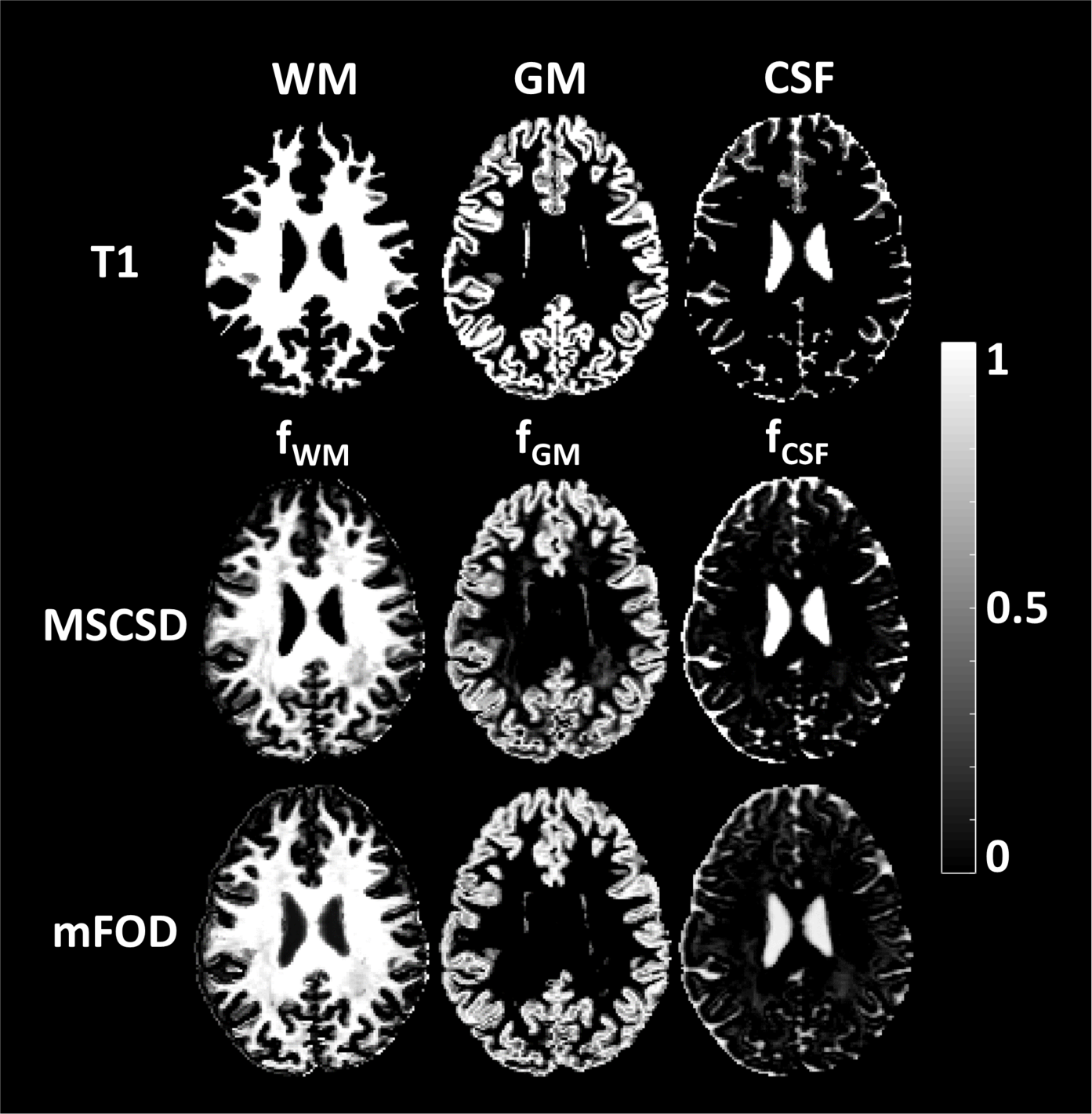
An example axial slice of the signal fractions estimated with the MSCSD and mFOD methods in correspondence of different response functions as compared to the tissue segmentation obtained from the structural T1 image. Estimates with the MSCSD and mFOD approaches are remarkably similar and have high anatomical correspondence with the tissue types that they are designed to represent.

#### FODs

The FODs estimated with the MSCSD and mFOD methods in two cortical areas are shown in Fig. 6 and Fig. 7 for an example axial slice and an example coronal slice, respectively. The WM fiber orientations estimated with the mFOD method (mFOD_1_) show remarkable similarity to the orientations estimated with MSCSD. Both FODs have very sharp and comparable orientations in the vicinity of WM and tend to become smaller towards deeper cortical GM. In contrast, the GM fiber orientations with the mFOD method (mFOD_2_) show dominant and anatomically plausible peaks in most GM voxel, with orientations mostly perpendicular to that of the underlying WM. When mFOD_1_ and mFOD_2_ are linearly combined (mFOD-WS), a continuous vector field from WM to deep cortical GM is revealed.

**Figure 6:**
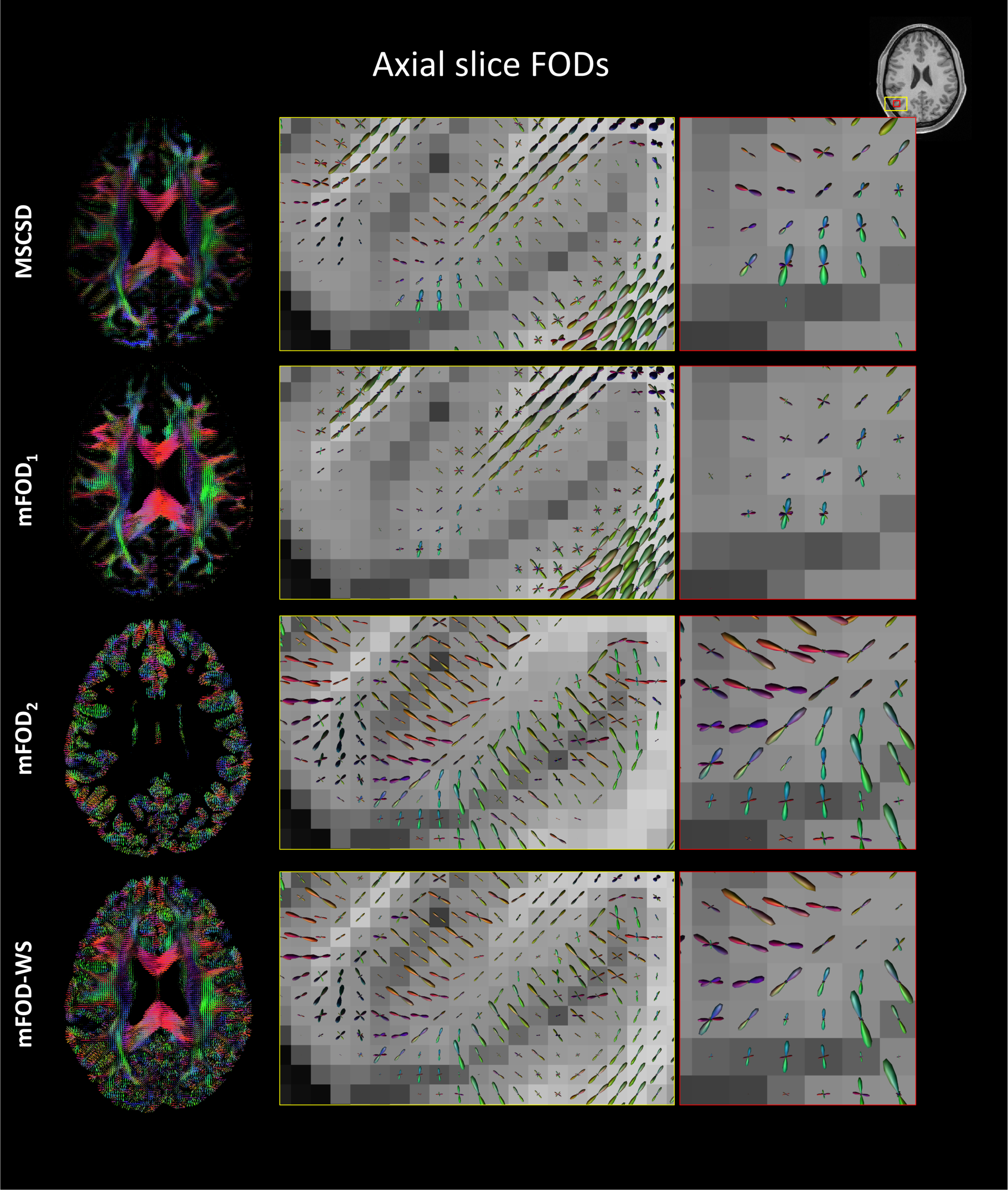
Fiber orientation distributions (FODs) obtained with MSCSD and the mFOD method in an axial slice. The middle and the right columns show increasing zoom levels to highlight details of FODs. The mFOD_1_ and mFOD_2_ refer to the FODs computed when using response functions meant to represent WM and GM, respectively. The mFOD-WS is the sum of mFOD_1_ and mFOD_2_ weighted by their corresponding signal fractions. The mFOD_2_ is non-zero only in GM and its proximity, and shows dominant orientations also in the deeper cortex, in contrast to both MSCSD and mFOD_1_. The mFOD-WS combines the information content of mFOD_1_ in WM with that of mFOD_2_ in GM.

**Figure 7:**
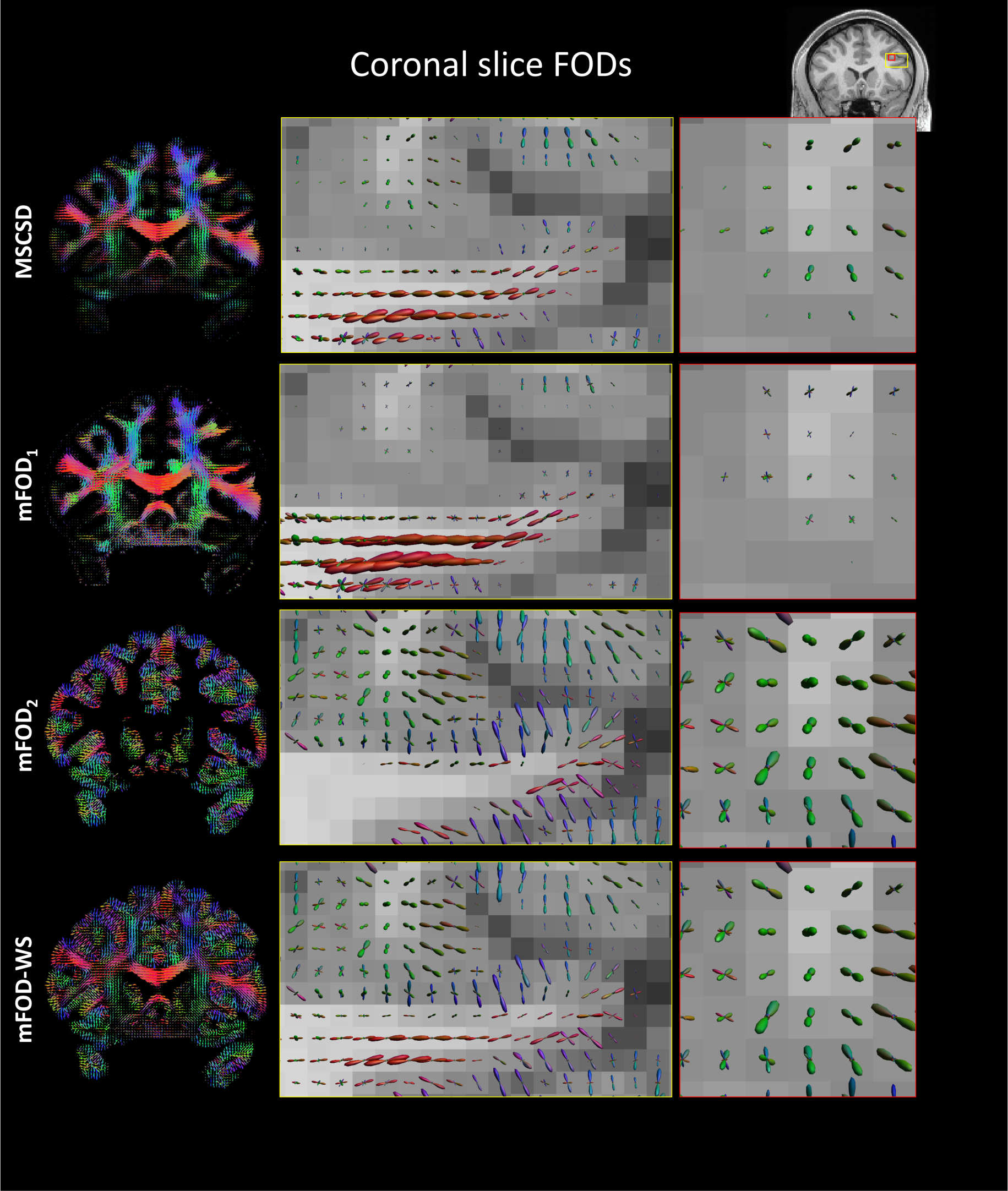
Fiber orientation distributions (FODs) obtained with MSCSD and the mFOD method in a coronal slice. The middle and the right columns show increasing zoom levels to highlight how the FODs change in vicinity of GM. The mFOD_1_ and mFOD_2_ refer to the FODs computed when using response functions meant to represent WM and GM, respectively. The mFOD_2_ is non-zero only in proximity of GM, and shows dominant orientations also in the deeper cortex, in contrast to both MSCSD and mFOD_1_. The mFOD_1_ provides directional information in strong agreement with MSCSD but occasionally shows more peaks.

Fig. 8 shows the FODs determined with mFOD-WS and MSCSD in an axial slice of the thalamus. The FODs derived with the two methods appear overall similar at visual inspection, but those determined with mFOD-WS appear sharper. A closer look (zoomed pane) reveals that the FODs computed with mFOD-WS have less angular uncertainty than those derived with MSCSD, and a better resolution of the peaks can be observed (white arrows). In the ROI shown in Fig. 8, the average number of peaks detected with MSCSD is 2.31 ± 0.88 as compared to 3.54 ± 1.03 with mFOD-WS. The average angle deviation between the main direction derived with mFOD-WS and MSCSD is 4.7 ± 4.5°.

**Fig. 8:**
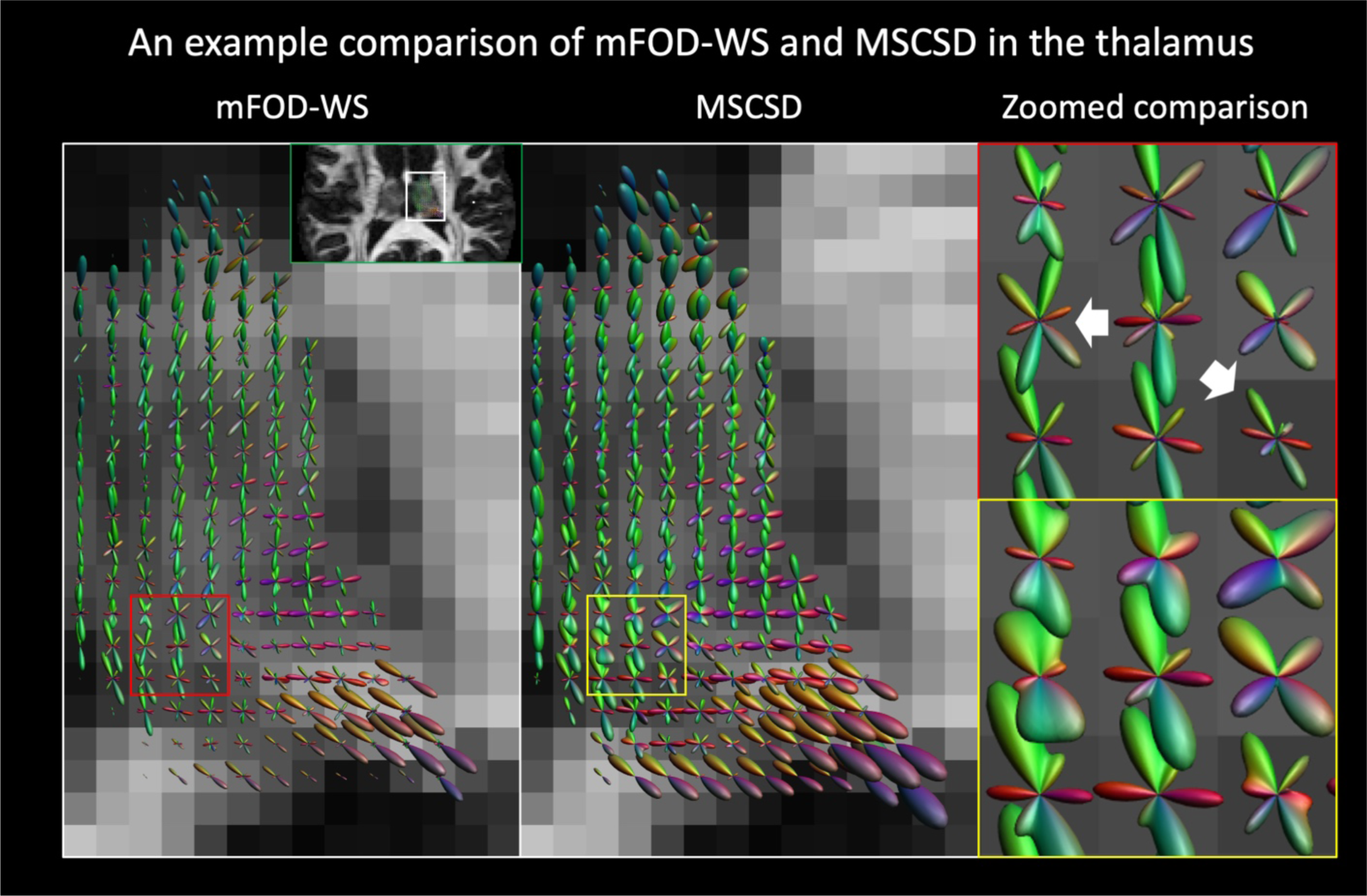
Fiber orientation distributions (FODs) obtained with the mFOD method and MSCSD in an axial slice of the thalamus. The right column zooms in a 3×3 neighborhood to better compare the visualized FODs. In general, the FODs in mFOD-WS appear sharper in the whole thalamus as compared to MSCSD. When looking into the zoomed region, the FODs in mFOD-WS appear sharper and appear to resolve complex configurations better than MSCSD (white arrows).

Fig. 9 shows the histogram of the peak deviations observed between MSCSD and mFOD-WS in WM, GM and at the WM/interface, as well as the angle between the FODs and the normal to the cortical folding as derived from the T1-weighted image with a structural tensor approach. When comparing the first peak direction of MSCSD and mFOD-WS in WM, we observe that more than 70% of the WM voxels have an angular deviation smaller or equal than 10 degrees. For the second and third peak we observed similar results, but also the presence of clusters of voxels with angle differences equal to 55° and 90° between the two methods. In GM and at the WM/GM interface, the angular deviation between the two methods is larger, up to 29 degrees, and the peak of the distributions are centered at about 0.1 degrees and 5 degrees, respectively. When looking at the angle with the cortex normal, the histogram of both peaks is centered at 20 degrees, but the mFOD-WS method results on average in a smaller angular deviation than MSCSD.

**Figure 9:**
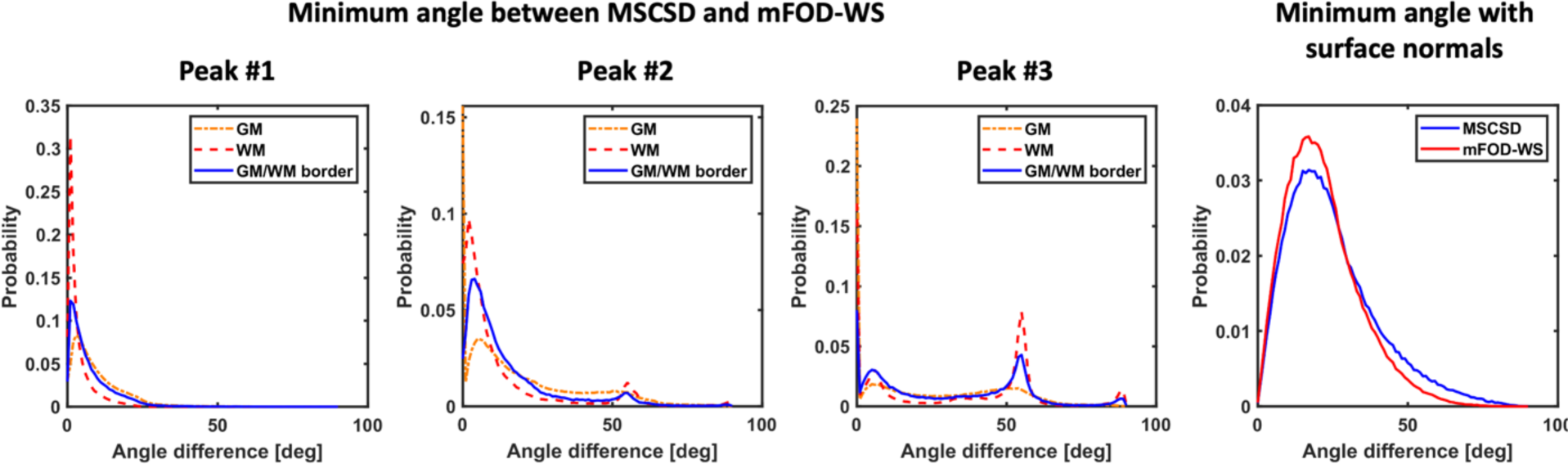
The first three histograms show the normalized frequency (probability) of the angle between each mFOD-WS peak and the closest MSCSD peak in WM, GM and at the WM/GM interface, respectively. The right histogram shows the probability of the angle difference between the normal direction to the GM/WM surface and the main direction with MSCSD or mFOD-WS. With regards to the left plots, the first peak of two methods has very small angle differences in WM, within 10 degrees in over 70% of voxels, whereas larger differences are observed in GM and at the WM/GM border. Larger differences were observed for the second and the third peak, with a subset of the voxels showing an angle of 60 degrees between the two methods. mFOD-WS resulted in lower angle deviation with the surface normal (derived with the structural tensor technique) as compared to MSCSD.

### Fiber tractography

Fiber tractography results in a frontal lobe region computed with the MSCSD and mFOD-WS methods are shown in Fig. 10. The fiber pathways are color encoded according to the tissue type they are spatially traversing, and show that the mFOD-WS method generates more dense tractography results in GM. Further, the fiber bundles reconstructed with the mFOD-WS approach cover the GM more uniformly and traverse the tissue with a more plausible shape and to a greater extent.

**Figure 10:**
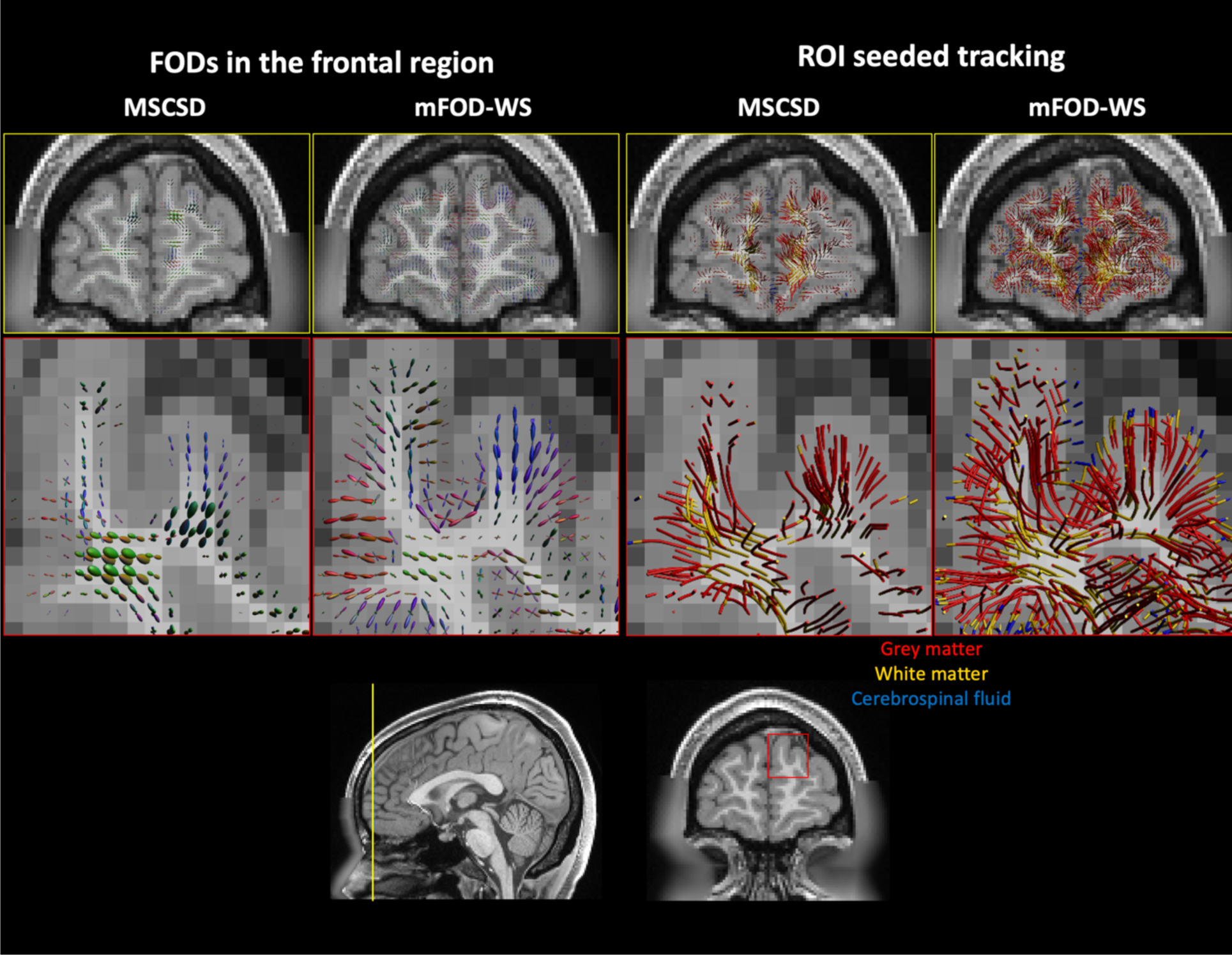
On the left, the FODs obtained with MSCSD and mFOD-WS in a frontal region. On the right, the fiber pathways obtained with the MSCSD and mFOD-WS methods by seeding in the shown axial slice (shown in yellow in the bottom left corner) with identical tracking parameters and FOD thresholds. The fiber pathways are color encoded according to the tissue type they transverse, as determined on the structural T1 image. The tracking with the mFOD-WS method resulted in a more complete tracking of the frontal cortex as compared to MSCSD, with the reconstructed trajectories covering a larger distance within the cortex.

Subsequently, we performed the whole-brain fiber tractography with the MSCSD and mFOD methods, both individually (mFOD_1_, mFOD_2_) and after the weighted sum procedure (mFOD-WS). The tracking with mFOD_1_ produces a whole brain reconstruction that is largely comparable to that of MSCSD, suggesting preservation of the WM tracking performance. Conversely, the tracking of mFOD_2_ results mostly in cortical reconstructions and short U-fibers, in-line with the GM-specific nature of this FOD. For pathways reconstructed with the mFOD-WS approach, 42.1% have their end points in GM, as compared to 34.1% when performing tractography with MSCSD. This is qualitatively visible in the tissue-type encoded reconstruction in Fig. 11, which shows that trajectories reconstructed with the mFOD-WS method cover the GM more extensively as compared to MSCSD. Note, however, that this also results in introducing extra spurious fibers.

**Figure 11.**
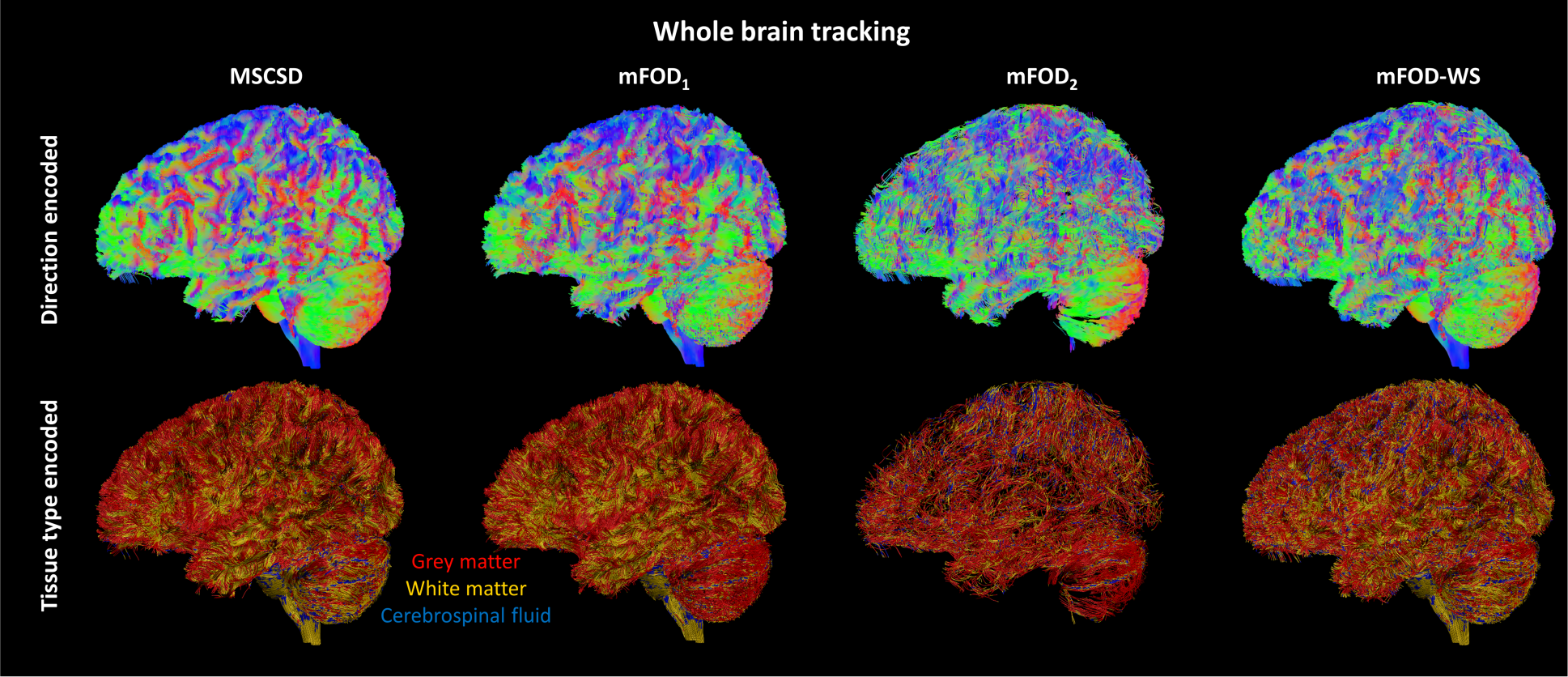
Whole-brain tracking with MSCSD and the mFOD methods. The first row shows the fiber pathways color encoded with the conventional color scheme, whereas the second row shows the same tracking color encoded by traversed tissue type (subsampled with factor 8). MSCSD and mFOD_1_ provide similar tracking results with the latter showing better characterization of the cerebellar cortex, as shown by the more homogeneous red coloring in the tissue type encoded tractography. The tracking with mFOD_2_ resulted exclusively in cortical and short (U-fibers) connections. With mFOD-WS, a more complete coverage of the GM is achieved as compared to both mFOD_1_ and MSCSD.

An example sagittal slab of the trajectories reconstructed with MSCSD and the mFOD-WS approach was obtained by cropping the whole brain fiber tractography, as shown in Fig. 12. In the first column, the trajectories are color encoded according to their distance from the WM/GM interface (blue) to the GM/CSF interface (red). It can be seen that the fiber bundles reconstructed with the mFOD-WS method consistently travel a longer distance within the cortical GM than with MSCSD, in-line with observations in Fig. 10. Further, the frontal, parietal and occipital side of the cortex appear to be better reconstructed with the mFOD-WS method.

**Figure 12.**
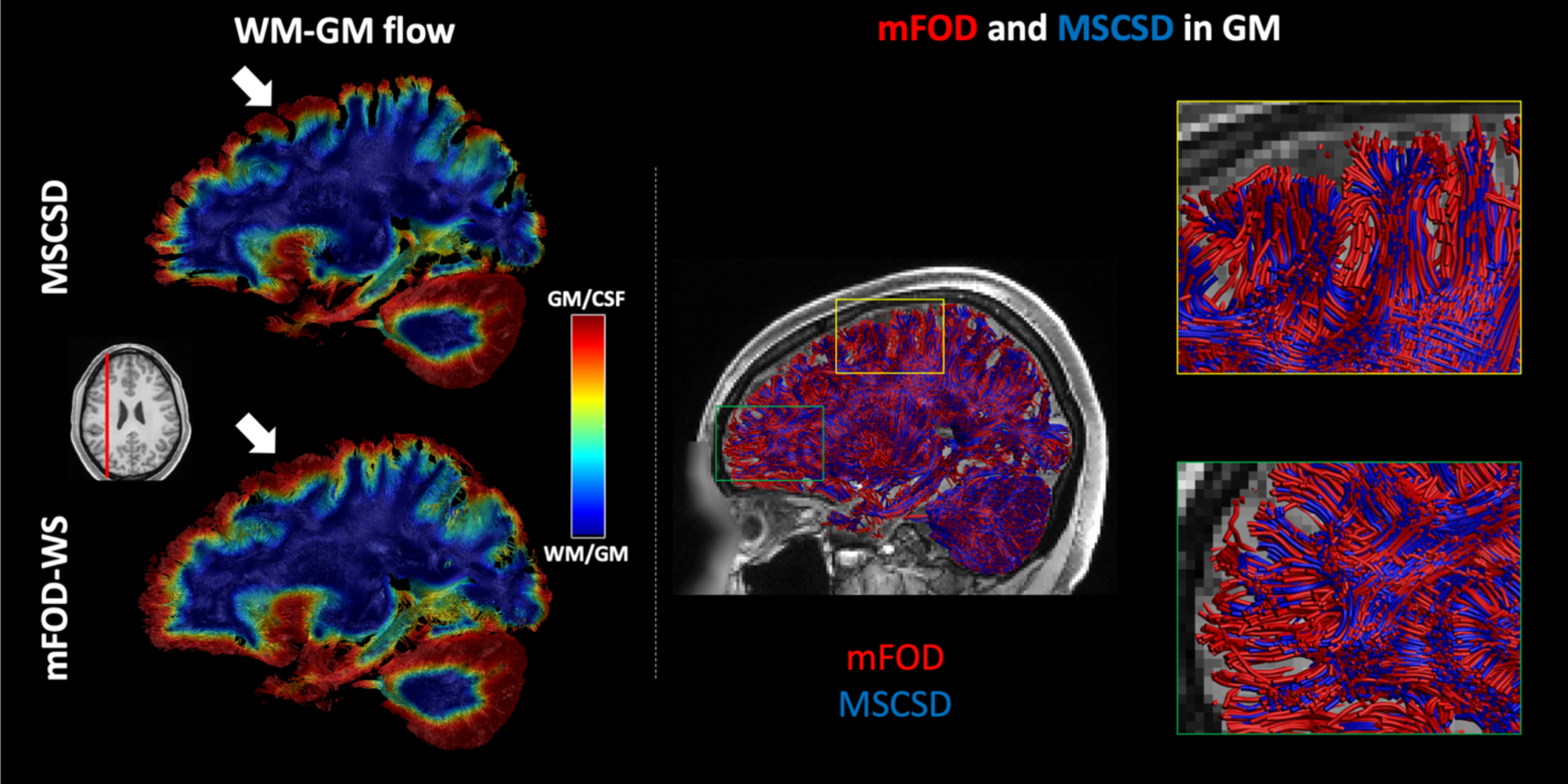
A sagittal slab (red line in the bottom left picture) of the fiber tractography results obtained with MSCSD and the mFOD-WS method. The first column shows the fiber pathways color encoded according to their normalized distance from the GM/WM and the GM/CSF interfaces. The distance was determined in 3D as flow (linear interpolation field) between the two surfaces using the same method we previously used to determine the cortex normal. The central and right column show the same sagittal slab color encoded by reconstruction method, and its zoom in a cortical region. The WM-GM flow color encoding shows that a deeper cortical tracking was achieved with the mFOD-WS method as compared to MSCSD, especially in the frontal and occipital areas, as well as a more contiguous coverage of the cortex (white arrows). When looking at the tissue type encoded images, mFOD-WS resulted in a more extensive and contiguous reconstruction of the cortical folding, and the tracts travelled deeper into the cortical GM, although a larger number of spurious reconstructions can also be observed.

While the mFOD framework has been primarily developed to target the cortical GM, deep GM regions as the thalamus have a central role in brain connectivity. In Fig. 13, we investigated the application of fiber tractography with mFOD-WS and MSCSD using a seed region covering part of the thalamus in one single axial slice. The fiber tractography obtained with both methods reveal the existence of thalamic – temporal and thalamic-sensory motor connections. With mFOD-WS, however, we also observe a remarkably larger number of streamlines reaching the brainstem, and the occipital cortex. Additionally, mFOD-WS results in longer reconstructions of the fiber tracts from the thalamus to both the sensory-motor and the frontal/pre-frontal cortex, in agreement with previous population studies^57^.

**Figure 13:**
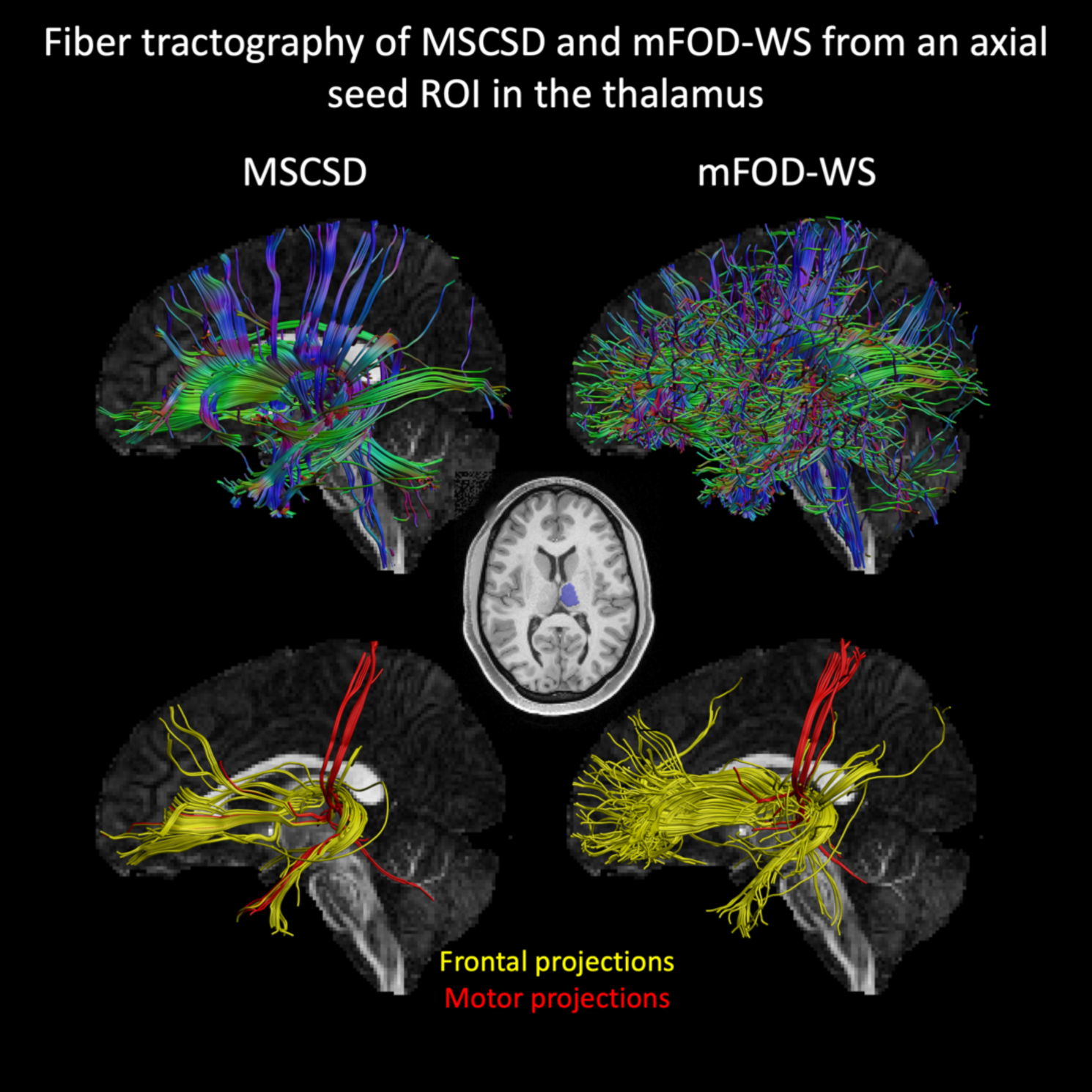
Results of fiber tractography of MSCSD and of the mFOD-WS method with a seed ROI located the Thalamus in one single axial slice. The streamlines reconstructed with mFOD-WS reached a larger extent of the cortex, especially in the occipital and frontal cortex. While the connection between the Thalamus and the sensory motor cortex was observed with both MSCSD and mFOD-WS, streamlines reconstructed with the latter branched more plausibly into the cortex (red tracts). When looking at the thalamic-frontal projection (yellow tracts), the fiber reconstructions with mFOD-WS covered a remarkably larger extent of the frontal cortex as compared to those obtained with MSCSD.

Fiber tractography of the mFOD-WS method using the HCP data is shown in Fig. 14 for different choices of tracking criteria. Good coverage of the cortical structure is observed for all settings, even when enforcing a minimum length of 50 mm, ensuring that most of the reconstructions originating in GM can reach the WM. When using an FOD threshold equal to 0.1, more spurious reconstructions are revealed as compared to a more restrictive threshold (0.3), but a denser tract reconstruction in the cortical ribbon and more extensive coverage of the contralateral hemisphere can be observed. Allowing for trajectories as short as 10 mm, results in a very dense reconstruction of the cortical gyri, showing short bundles that seem to travel tangentially to GM, as well as many short radial reconstructions.

**Figure 14.**
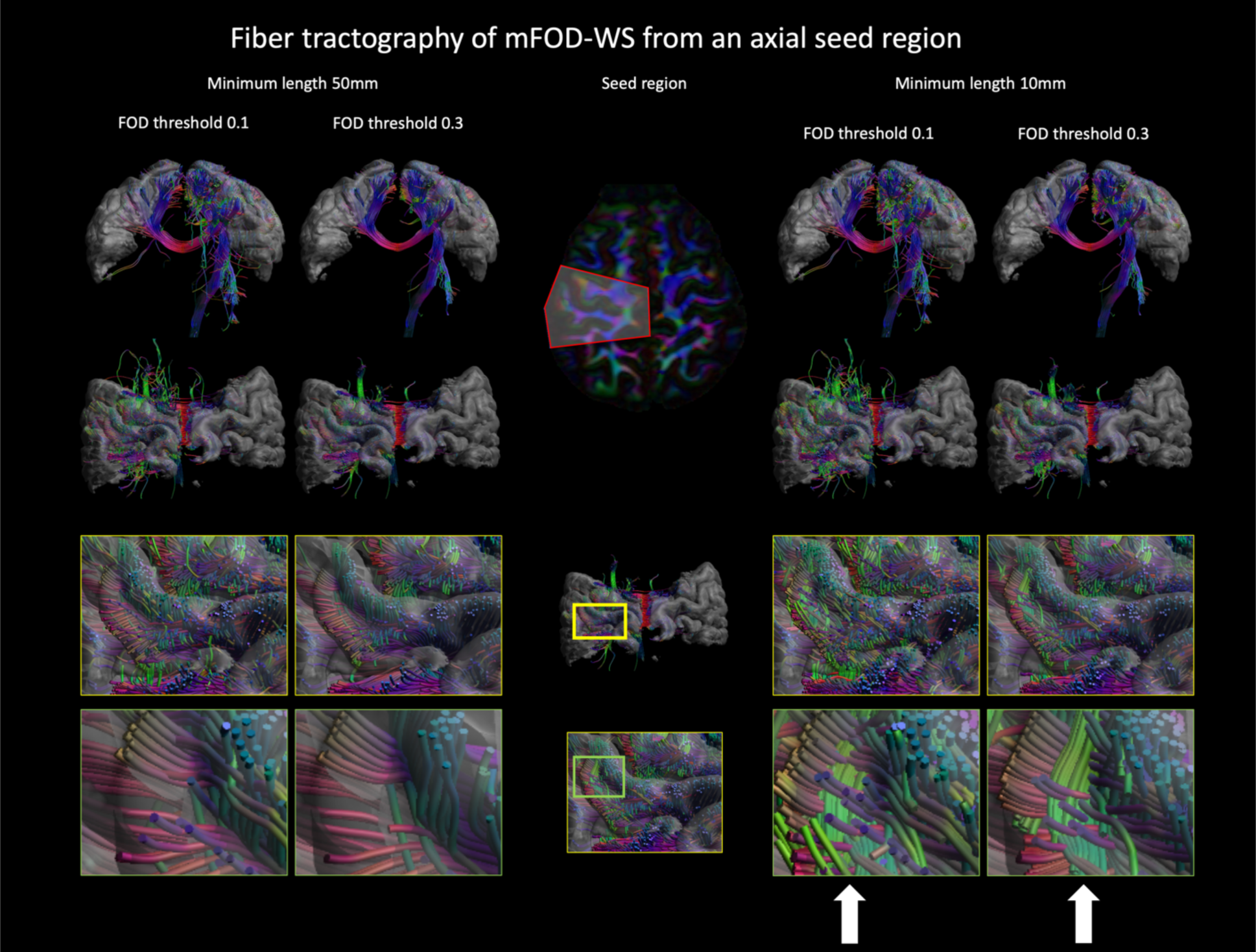
Results of fiber tractography of the mFOD-WS method with a seed ROI in an axial slice including the left central superior cortex. The tractography results are shown for two choices of FOD thresholds and minimum tract length. Reconstruction of WM bundles reaching the cortex can be observed with all tested settings. A lower FOD threshold (0.1) results in a more contiguous representation of the gyri as compared to the more restrictive setting (0.3), but also in more false positive reconstructions. Lowering the minimum tract length from 50 mm to 10 mm allows one to reconstruct short fibers tangent to the cortex (white arrows) that appear to travel within the cortical ribbon.

## Discussion

In this work, we have introduced the mFOD framework to perform spherical deconvolution of multiple anisotropic response functions. With simulations and HCP data, we have shown that it is possible to reconstruct two anisotropic FODs, specific to WM and GM, while also accounting for the CSF partial volume effect. Further, we have shown that the mFOD approach improves the reconstruction of FODs in GM as compared to existing state-of-the-art methods, and that this results in improved fiber tractography reconstruction in regions near or in the GM.

### Feasibility of mFOD

The reconstruction of multiple anisotropic FODs is a complicated, numerical challenge that requires the inversion of a matrix whose rank (number of independent rows/columns) is much lower than the number of possible solutions (FOD amplitudes on the unit sphere), a recurring challenge in deconvolution problems. Previous studies, as Tournier et al.^35^ and Jeurissen et al.^31^ used an L2 regularization term enforcing continuity (smoothness) in the FOD amplitudes. In this work, we used a non-negative least-squares solver with L2 regularization encapsulated in an iterative solver^44^. Taking into consideration that the mFOD formulation (eq. [2]) hosts multiple FODs, this might be suboptimal and introduce discontinuities in the solution. Nevertheless, our results suggest that the mFOD approach does not propagate smoothness between multiple FODs, likely due to the attenuation of small perturbations imposed by the median operator^43^. Indeed, the results of simulations, shown in Fig. 2, Fig. 3 and Fig. 4, suggest that this approach allows to effectively disentangle the partial volume effects between mixtures of isotropic (n=1) and anisotropic (n=2) signal mixtures, if sufficient SNR is provided. In particular, the partial volume effects between WM and CSF are the easiest to disentangle, and an SNR equal or greater than 20 is sufficient to obtain accurate separation of signal fractions and minimal angular error on the WM FODs. In contrast, the separation of GM from both WM and CSF is a harder task, probably as a result of the limited anisotropy of this tissue type. In Fig. 4, we observe that the GM estimation shows a small but consistent offset from the simulated values. This is very likely caused by the abovementioned median operator in the deconvolution framework (GRL) on which mFOD is built. Such operator is designed to be used in combination with deconvolution matrices containing highly anisotropic signal basis. When spherical deconvolution is performed with a deconvolution matrix including moderately anisotropic elements (e.g. generated with the NODDI model with *κ* = 1), this is likely to produce some genuine small amplitudes which are discarded by the median filtering, causing a small but consistent bias in the f_GM_ estimate. Of note, in our simulations we did not compare the performance of mFOD to MSCSD, because such method is data-driven, e.g. it estimates its deconvolution matrix from the data, which is not feasible in a single-voxel simulation. Furthermore, we believe that such comparison would not be completely fair, as by design the MSCSD method lacks the ability to fit the anisotropic GM-like signal that we introduce in the simulated signals.

If an SNR equal or above 50 is achieved, the mFOD framework can preserve the ratios between different tissue types and can estimate their mixtures with minimal biases (Fig. 2 and Fig. 3). While an SNR of about 50 might be difficult to achieve in clinical settings, accurate pre-processing and denoising^58^, or even deep-learning based reconstructions^59^ might mitigate this need. According to Fig. 3, the crossing angle between a WM bundle entering into the cortex and the cortical folding itself is likely to have an impact on the performance of mFOD. Generally, white matter bundles travel to their cortical endpoint perpendicularly^60^ to the cortical folding, with exceptions such as the primary visual cortex^20^ or the U-fibers adjacent to the cortex^14^. If the direction of the axons leaving the cortex (to the WM) would not match that of intra-cortical axons, the results of the simulations suggest that the GM FOD would be estimated with an angular error between 15 (SNR 30) and 10 degrees (SNR 50), which are likely to perturbate the tracking trajectory. In practice, very high SNR is likely required to reconstruct fiber pathways tangential to the cortical folding, but less critical to detect radial trajectories. In light of the insights provided by simulation I, we conclude that mFOD can estimate the GM FOD with reasonable angular error in the cortex itself (configuration V in Fig. 3) and in the case of limited partial volume with a single WM fiber (configuration IV in Fig. 3). The performance of mFOD at separating WM-like and GM-like FODs is also confirmed by the in-vivo results, as the signal fraction map associated to GM is zero-valued in all non-GM voxels and has values of 1 in almost all GM voxels, which suggests excellent separation of the two tissue types. In proximity of the WM/GM interface, however, the possible presence of an additional crossing with the superficial U-fibers is likely to generate unreliable estimates (configuration III of Fig. 3), suggesting that further research is needed to address the challenge of crossing the superficial WM^13^. On the other hand, it should be noted that the above result corresponds to a GM-like signal generated with the NODDI parameter with *κ* = 1, which results in low signal anisotropy. While such value seems appropriate in the cortex, i.e., when f_GM_ = 1, at the WM/GM interface, higher values of *κ* are typically observed. According to Supplementary Figure S2, with a higher value of *κ*, the angular error of the GM FOD drops drastically.”

### GM modelling

While recent work has introduced the reconstruction of multiple WM FODs^33^ to account for inadequacies in WM modelling in infants, the mFOD framework has the unique characteristic of reconstructing multiple FODs that are specific of remarkably different tissue types as WM and GM, as shown in Fig. 6 and Fig. 7. The figure shows that WM fiber orientations estimated with mFOD (mFOD_1_) and MSCSD are very similar in WM, and that both exhibit very small values in GM. In contrast, the GM fiber orientations estimated with mFOD_2_ show consistent and well-defined diffusion directions for a large extent of the cortical GM. The performance improvement over MSCSD of the mFOD framework in GM is explained by considering the improved tissue modelling. Approaches as MSCSD essentially describe the WM and GM mixture as the sum of an anisotropic and an isotropic signal, similarly to the ball-and-stick model^61^, where the anisotropic signal is modelled with properties extracted from single fiber population WM^35,50,62^. Therefore, in practice, by applying any constrained spherical deconvolution method in GM, one assumes the axonal component of GM to produce a similar signal to that of a WM bundle without crossing fibers. However, assuming the cortical axons and dendrites of the cortex to be identical to the axons in the corpus callosum or other major WM bundles seems a stretch given their different diffusion signature^63^. In this work, we represent the GM signal with a special case of the NODDI model^38^, which represents a first approach to take into account additional effects as tortuosity^64^ and intra-to-extra neurite signal ratios into the deconvolution problem.

### How to use multiple FODs

One of the open challenges in the mFOD framework regards how to combine the multiple FODs computed by the method. In this work, we adopted a pragmatic approach and considered their weighted sum (mFOD-WS). This choice produces convincing and contiguous FODs, as shown in Fig. 6 and Fig. 7. However, this might be sub-optimal for fiber tracking approaches, as it might blur the peaks detected in individual FODs. Future work is needed to establish whether dynamically tracking the locally larger FOD, or dynamically choosing the FODs most aligned to the tracking direction might represent better alternatives. Additionally, sparsity constraints as L1 or L0 norm^65^ could be taken into consideration to improve the FOD separation and minimize overfit. Nevertheless, when we compared the main peaks of mFOD-WS to those obtained with MSCSD (Fig. 9), we observed remarkable similarity between the two in WM. Further, the average angular error between the two techniques was below 10 degrees in over 70% of WM voxels, suggesting that the mFOD-WS approach does not affect the performance of spherical deconvolution in WM. In GM, the mFOD-WS approach resulted in smaller angular deviations with respect to the cortex normal as compared to MSCSD. While the cortex normal might not always be representative of the direction with which axons are organized in the cortex, they represent an accepted assumption^18^, and might serve as a comparison term between the two methods. The average angular deviation between the cortex normal and mFOD-WS is 20 degrees, which is in-line with the angular error predicted in case of WM and GM mixtures at SNR 50 (Fig. 3).

### Improved fiber tractography in GM

Fiber tractography is one of the main applications of spherical deconvolution techniques. Here, we used a deterministic approach to facilitate the interpretation of pathway reconstructions. As shown in Fig. 10, we obtained a higher percentage of tract pathways reaching a GM region with the mFOD-WS approach than with MSCSD given identical tracking parameters. Furthermore, the fiber pathways reconstructed with the mFOD-WS method covered the cortical structure more extensively, and their geometrical configuration appeared more similar to previous reports of high resolution cortical dMRI^30^. While a similar tractography result might also be achievable with MSCSD by tuning the FOD threshold^66^, this is likely to introduce not only false positives in WM, but also to worsen the tracking in GM itself due to a larger number of spurious peaks^67^. One important application of fiber tractography is the study of deep GM structures, due to their central role as hubs for many brain connections. In Fig. 8, we investigated the performance of mFOD-WS in the thalamus. We observe that the use of mFOD-WS seems also beneficial in deep GM, as it provides FOD estimates that are sharper and better resolved than those provided by MSCSD, which is desirable in fiber tractography applications. Furthermore, in the thalamus, we found a small but consistent angular bias of about 5° between the main directions estimated with the two methods, and a higher number of detected peaks with mFOD-WS, 3.5 vs 2.3. When performing fiber tractography from the thalamus (Fig. 13), the fiber tracts reconstructed with mFOD-WS covered a larger extent of the brain as compared to the fiber tracts reconstructed with MSCSD, including previously reported connections to the occipital, sensory-motor and frontal cortices, although a larger number of spurious fibers was also observed. Furthermore, the reconstruction of the thalamic-frontal tract with mFOD-WS appears more in line with previous reports.

When performing whole-brain tractography, the use of the mFOD-WS approach resulted in more tract pathways having a start or endpoint in the GM (+8%), and generally in more extensive GM coverage, as shown in Fig. 11. This was even more apparent when looking at a sagittal slab (Fig. 12), which showed how fiber bundles could be followed in GM up to the GM/CSF interface, at the price of potentially more spurious reconstructions as compared to MSCSD. In practice, mFOD could be beneficial to applications such as pre-surgical planning^17^ or structural connectivity studies^68^, where propagating fiber pathways from the deep WM into the cortex is crucial. Existing fiber tractography methods mostly do not define the seed points in the cortex, but rather at the GM/WM interface or in the adjacent WM, potentially introducing a number of false positives^69^. For similar reasons, connectivity studies are often performed by first seeding WM or the WM/GM interface, then filtering the trajectories with anatomical constraints^70^, and finally assigning the fiber bundles to the closest GM node after spatial dilation, potentially introducing uncertainty in the assignment^71^. To overcome these limitations and reconstruct complete fiber pathways to/from the cortical start/endpoints, 4 major steps are required: i) the estimation of reliable FODs in WM, ii) crossing the superficial WM at the WM/GM interface, iii) overcoming the gyral bias in fiber tractography and iv) the estimation of reliable FODs in GM. The mFOD framework is a first step towards the solution of this comprehensive challenge, as it allows to robustly determine the FODs in cortical areas. This problem has also been investigated in previous studies and has been addressed with computational approaches based on the geometry of the cortex itself. While such methods have the advantage of not requiring elaborate diffusion acquisition schemes as mFOD, they simplify the structure of the cortex, enforcing fiber tractography to proceed radially to the cortex, which does not take into account the potential presence of tangential fibers or the adverse effects related to disease or abnormal development of the cortex organization. Taking into account the results shown in Fig. 14, it seems possible with mFOD to also effectively reconstruct some crossing bundles in the GM itself, where radial projections cross previously reported tangential fibers^20,21,26^. Fig. 14 also highlights the importance of the FOD threshold parameter in fiber tractography. When the FOD threshold is lowered, a larger number of fiber tracts is reconstructed. These fiber reconstructions are also generally longer and can become erratic when the FOD is not well-defined, such as in the outer cortex where the partial volume with CSF dominates the signal. Increasing the value of the FOD threshold reduces the number of spurious fibers, but likely also the number of genuine pathways that are discarded^69^. The accurate estimation of FODs at the WM/GM interface is more challenging than for GM itself according to the results of simulation I (Fig. 3). In our in-vivo experiments with mFOD, fiber tractography generally reached the cortex, but this might be more challenging in datasets with lower SNR. In that case, the mFOD approach might be further enhanced by taking into account geometric frameworks such as the “Surface Enhanced Tractography”^19^ or the “Structure Tensor Informed Fiber Tractography”^54^, or with hybrid acquisition strategies aimed to better resolve the FODs at the WM/GM interface^72^. In this manuscript, we have employed a deterministic tractography method to focus on the FOD estimation step. However, mFOD could also be used in combination with probabilistic fiber tractography^73^, multi-peak tractography^74^ or even global tractography^75^, as shown in Supplementary Material Figure S4.

### Methodological considerations

The models used here to describe WM and GM properties were chosen from current popular methods, but their combination might not be the optimal representations to estimate multiple FODs. The DKI model, here used in its simplified version with isotropic kurtosis^47^, efficiently represents non-Gaussian signals with a limited number of parameters but is sub-optimal to account for direction dependent signal restrictions. Alternatively, geometric representations of the WM signal^49^, as well as data driven methods^50^ could also be explored. Note that the diffusion approaches suggested for WM (DKI) and GM (NODDI) could be further adjusted or even replaced by other more specific tissue models to further improve the mFOD framework. In name of clarity, we fixed the parameters of the NODDI model to the average values observed in GM, but these values could be suboptimal at voxel level. In Supplementary Figure S1, we have briefly investigated the impact of mis-estimating the NODDI parameters in mFOD, observing that overestimating the value of *κ* in mFOD results in worse angular errors in the GM FOD but minimal or no effects on the signal fraction estimates. Conversely, underestimating the value of *κ* has marginal effects on the angular error, but biases the signal fraction estimates.

The dMRI acquisition protocol is expected to have a major influence on the performance of the mFOD framework. Here, however, we aimed to show a proof-of-concept of the feasibility of the mFOD approach, and therefore relied on the high-quality HCP data. Nevertheless, in the Supplementary Material we briefly investigated with simulations the impact on the performance of mFOD of specific changes to the acquisition protocol. Supplementary Figure S3 shows that halving the acquisition protocol with respect to the HCP gradient schemes would minimally affect the estimation of the WM FOD and of the signal fractions, but would worsen the precision of the estimates, as expected. The reduced acquisition protocol negatively impacts also the angular error of the GM FOD, increasing the error of about 5° for the 90° WM-GM crossing simulation. In Supplementary Figure S2, we investigated how different distributions of HCP-like multi-shell diffusion weightings would impact the mFOD performance. We observe that acquiring more directions in the outer shell or reducing its diffusion weighting to b = 2500s/mm^2^ does not improve nor worsen the angular error of the GM FODs. Conversely, increasing the maximum diffusion weighting to b = 4000 s/mm^2^ seems to improve the estimation of the GM FOD, in line with recent reports using ultra-high diffusion weightings^39,76^.

In Fig. 9, we used the cortex normal direction as a comparison for FODs computed with both MSCSD and mFOD. While it is reasonable to assume that most axonal projections will traverse the cortex with a perpendicular trajectory, this is not always the case. Furthermore, the method employed to compute the normal trajectory is not accurate when the sulci are separated by less than 2 voxels and should therefore be interpreted cautiously. The performance of the mFOD framework has been shown for two subjects and, although promising, should be further evaluated on larger cohorts and with more clinically achievable protocols. Finally, while we have shown that the mFOD framework improves the performance of spherical deconvolution and fiber tractography in GM, it remains to be shown to which extent such improvements will benefit clinical studies and structural connectivity investigations.

In conclusion, we have presented the mFOD framework to perform tissue-specific spherical deconvolution which can simultaneously reconstruct multiple distributions of fiber orientations per voxel. By integrating specific WM and GM models, we have shown that the mFOD approach can improve the performance of spherical deconvolution in GM while retaining the performance of state-of-the-art methods designed for WM.

## Acknowledgements

The research of A.D.L. is supported by the ERA-NET Neuron grant R.4195 – “Repetitive Subconcussive Head Impacts - Brain Alterations and Clinical Consequences” (REPIMPACT). The research of F.G. is funded by the Chinese Scholarship Council (CSC), No. 201306080017. The research of A.L. is supported by VIDI Grant 639.072.411 from the Netherlands Organisation for Scientific Research (NWO).

