## Supplementary Material for "Spherical deconvolution with tissue-specific response functions and multi-shell diffusion MRI to estimate multiple fiber orientation distributions (mFODs)"

### Additional simulations

In the following sections, we report the results of additional simulations based on the methodology used for Simulation I in the main text.

#### Effect of the NODDI $\kappa$ parameters on mFOD

In this section, we investigate the effect on the mFOD performance of the NODDI parameter $\kappa$ imposed in the signal simulations ($\kappa_{I}$), and of the parameter $\kappa$ used to generate the deconvolution matrix H_GM_ ($\kappa_{E}$). Following the same procedure of Simulation I in the main text, we simulated a 45° fiber crossing of a WM-like and a GM-like fibers with equal signal fraction (0.5). The simulation was repeated 1000 times by adding Rician noise at SNR levels in the range 10-150 for five different settings of the NODDI parameter $\kappa:$ i) $\kappa_{I}=\kappa_{E}=1$, ii) $\kappa_{I}=\kappa_{E}=2$, iii) $\kappa_{I}=\kappa_{E}=3$, iv) $\kappa_{I}=2, \kappa_{E}=1$, v) $\kappa_{I}=1, \kappa_{E}=2$.

The results of the simulation, reported in Figure S1, show that the angular error of the GM FOD decreases with increasing value of $\kappa_{I}$. This is expected, as higher values of $\kappa_{I}$ imply lower axonal dispersion and thus lower uncertainty on the peak direction of the corresponding FOD. The signal fraction error and the angular error are both minimal when the correct value of $\kappa_{E}$ is used, e.g. $\kappa_{E}=\kappa_{I}$. A lower value of $\kappa_{E}$ than $\kappa_{I}$ results in biases in the signal fraction but does not seem to worsen the angular error. Conversely, a value of $\kappa_{E}$ larger than $\kappa_{I}$ introduces in the corresponding FOD orientation, but not in the signal fraction estimates.

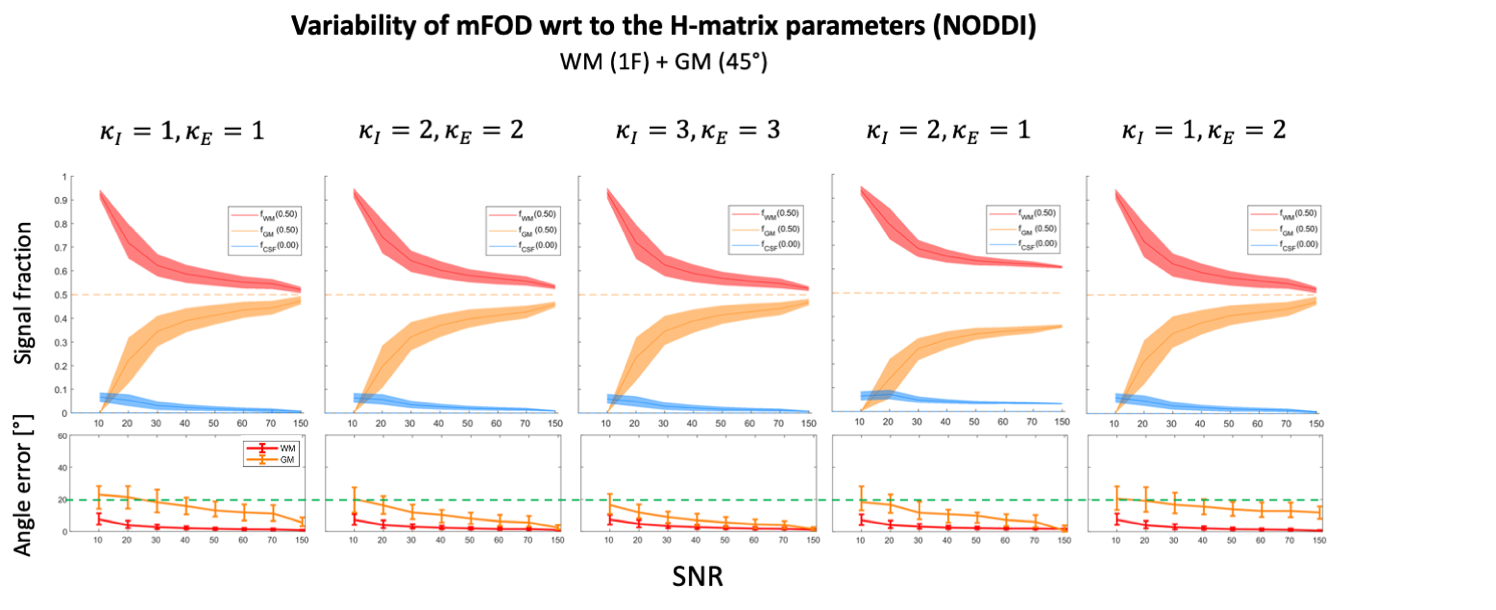

Figure S1: Simulations of a WM-like and a GM-like crossing fibers with equal signal fraction and crossing angle 45°. For each simulation, the top plot shows the 25th – 75th percentile (shaded area) and the median (solid line) of the signal fractions estimated with the mFOD approach for different SNR levels. The bottom plot shows the angular error of the main direction of the WM and GM FODs as compared to the ground truth value. The solid line represents the median value, whereas the error bars the 25th and 75th percentile. The green dotted line corresponds to an angular error equal to 20°.

#### Effect of the acquisition protocol on mFOD

In this section, we investigate the impact of the diffusion weighting scheme on the mFOD performance with simulations based on Simulation I. We focus this analysis on the HCP multi-shell gradient scheme, which has been used throughout this work as baseline, and investigate whether acquiring a larger amount of gradient directions in the largest shell (to compensate the lower effective SNR), or using a lower / higher maximum diffusion weighting affects the performance of mFOD. To this end, we simulated the crossing of a WM-like and a GM-like fibers with crossing-angle 90°, with I) the HCP-like protocol, II) with a 3 shell scheme including 30 volumes at b = 1000 s/mm^2^, 30 volumes at b = 2000 s/mm^2^, and 210 volumes at b = 3000 s/mm^2^, and with the HCP-like protocol but maximum diffusion weighting III) b = 2500 s/mm^2^ and IV) b = 4000 s/mm^2^. Additionally, the same simulation was repeated with an acquisition protocol including only half of the directions of the original HCP protocol for crossing angles 10°, 45° and 90°.

Figure S2 shows that shifting more directions to the outer shell slightly reduces the angular error of the WM FOD (-8% at SNR 50) as compared to the HCP-like protocol, but it does not reduce the GM FOD angular error (-0.05% at SNR 50) and it slightly worsens the signal fraction estimates (at SNR 50, +1.24% for f_WM_, +2% for f_GM_). Reducing the diffusion weighting of the outer shell to b = 2500 s/mm^2^ worsens the angular error of both the WM FOD and the GM FOD, as well as the signal fraction estimates as compared to the HCP protocol. Conversely, increasing its maximum diffusion weighting to b = 4000 s/mm^2^ reduces the angular error of both the WM (-23% at SNR 50) and the GM FODs (-21% at SNR 50), and improves the precision of the signal fraction estimates.

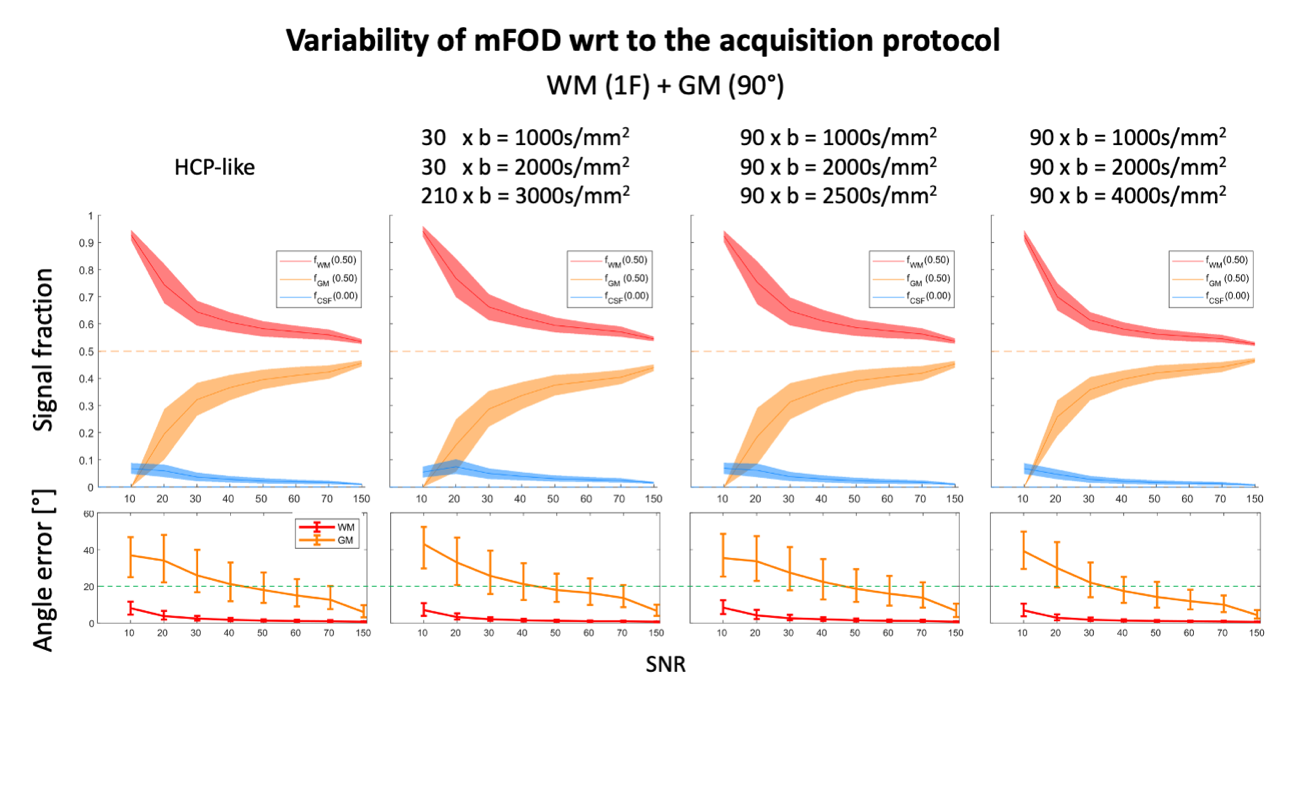

Figure S2: Simulations of a WM-like and a GM-like crossing fibers with equal volume fraction and crossing angle 90°. For each simulation, the top plot shows the 25th – 75th percentile (shaded area) and the median (solid line) of the signal fractions estimated with the mFOD approach for different SNR levels. The bottom plot shows the angular error of the main direction of the WM and GM FODs as compared to the ground truth value. The solid line represents the median value, whereas the error bars the 25th and 75th percentile. The green dotted line corresponds to an angular error equal to 20°.

Figure S3 shows the results obtained when applying mFOD to signals simulated with the HCP protocol as compared to a reduced protocol including only half of the original directions. Reducing the number of acquired gradient directions leads to larger errors in the signal fraction estimates. Further, it worsens their precision and increases the angular error of the GM FOD. This is expected, as reducing the number of acquired gradient directions essentially reduces both the angular resolution of the data as well as its effective SNR.

Taken together, the results of Figure S2 and Figure S3 suggest that mFOD requires a considerable amount of data to achieve a high effective SNR, in order to produce reliable estimates. Furthermore, the use of a maximum diffusion weighting larger than b = 3000 s/mm^2^ seems advisable and should be investigated in future studies.

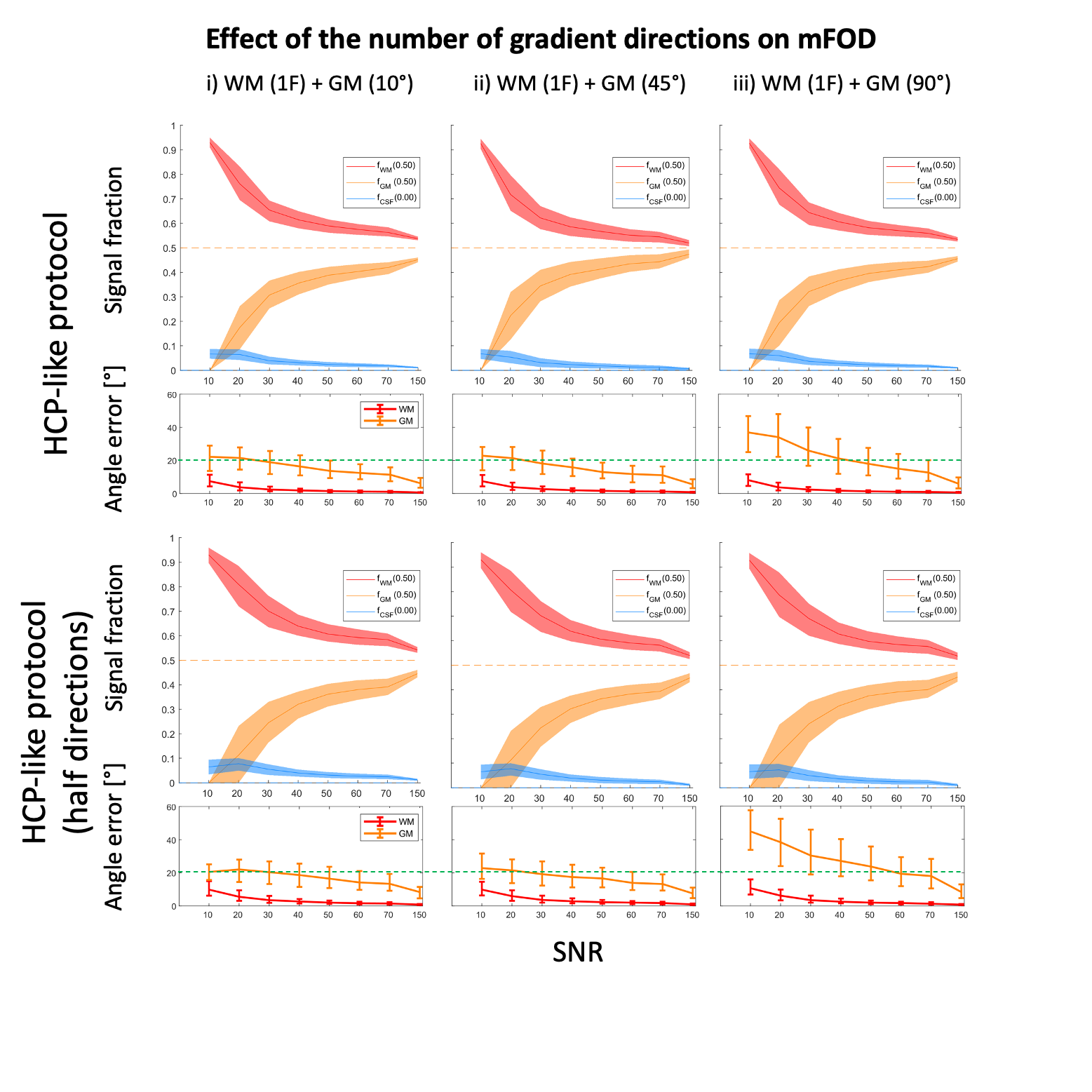

Figure S3: Simulations of a WM-like and a GM-like crossing fibers with equal volume fraction and crossing angles 10°, 45°, 90°. For each simulation, the top plot shows the 25th – 75th percentile (shaded area) and the median (solid line) of the signal fractions estimated with the mFOD approach for different SNR levels. The bottom plot shows the angular error of the main direction of the WM and GM FODs as compared to the ground truth value. The solid line represents the median value, whereas the error bars the 25th and 75th percentile. The green dotted line corresponds to an angular error equal to 20°.

### Probabilistic and multi-peak tractography of mFOD-WS

Fiber tractography with probabilistic and multi-peak approaches are often used as alternatives to deterministic tracking as they have been shown to provide more complete fiber reconstructions, although generally at the price of a larger number of false positive reconstructions. In this section, we qualitatively compare the application of deterministic, probabilistic and multi-peak fiber tractography to mFOD-WS and MSCSD in a small cortical region of the left parietal cortex. Deterministic fiber tractography was performed in ExploreDTI, seeding in a 2x2 voxel ROIs placed in the deep WM of the gyrus, with step-size 0.6mm, FOD threshold 0.1, angle threshold 45° and seed resolution 0.1 x 0.1 x 0.1mm^3^. Probabilistic tractography was performed in MRtrix 3 ([www.mrtrix.org](http://www.mrtrix.org)) using the iFOD2 approach, 1000 iterations, step-size 1mm, FOD threshold 0.1 and default angle threshold and seed resolution. Multi-peak tractography iteratively repeats deterministic tractography, and at each iteration it branches new streamlines from th existing along previously unused FOD peak directions. We performed the multi-peak tractography with custom code available as part of MRIToolkit (<https://delucaal.github.io/MRIToolkit/>), using the same parameters of deterministic tractography but with a lower seed resolution equal to 1.25x1.25x1.25mm^3^, for computational reasons.

The results obtained with the three tractography methods are reported in Figure S4. The figure shows that the fiber tracts reconstructed with mFOD-WS travelled deeper in the cortex and covered a larger extent of the grey matter as compared to those reconstructed with MSCSD, in line with the results shown in the main text. With deterministic tractography of mFOD-WS, all pathways enter the cortex with minimal curvature as compared to their trajectories in WM, a known limitation of fiber tractography that causes the well-known gyral bias. The streamlines reconstructed with probabilistic tractography of mFOD-WS cover a larger extent of the cortical grey matter as compared to deterministic tracking, but almost no streamline is observed entering the gyral banks. Different results are observed with the multi-peak tractography of mFOD-WS, as the streamlines cover a large extent of the gyral crown but also reach the gyral banks. However, the latter approach also produces a larger number of apparently spurious reconstructions. These results suggest that the tractography method of choice has a major impact on the reconstructed streamlines, and that future work should investigate whether the combination of mFOD with approaches other than deterministic tractography could be beneficial to improve the quality of the fiber reconstructions in GM.

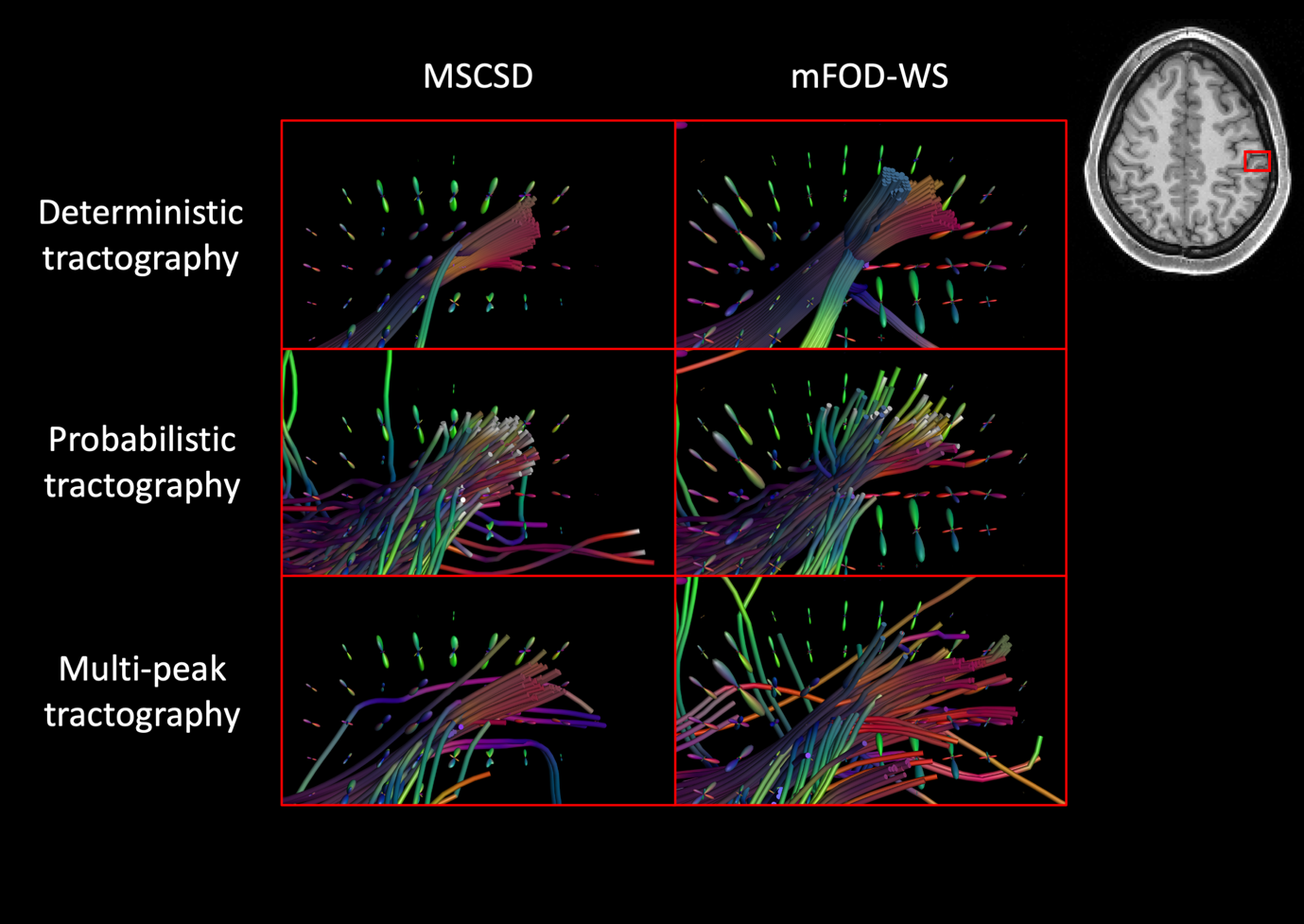

Figure S4: The results obtained when performing fiber tractography with MSCSD and mFOD-WS in the proximity a gyrus in the left parietal cortex with a deterministic, a probabilistic and a multi-peak approach.
